# Adiponectin and related C1q/TNF-related proteins bind selectively to anionic phospholipids and sphingolipids

**DOI:** 10.1101/776690

**Authors:** Jessica Ye, Xin Bian, Jaechul Lim, Ruslan Medzhitov

## Abstract

Adiponectin (also known as Acrp30) is a well-known adipokine associated with protection from cardiovascular disease, insulin resistance, and inflammation. Though multiple studies have investigated the mechanism of action of adiponectin and its relationship with tissue ceramide levels, several aspects of adiponectin biology remain unexplained, including its high circulating levels, tendency to oligomerize, and marked structural similarity to the opsonin C1q. Given the connection between adiponectin and ceramide metabolism, and the lipid-binding properties of C1q, we hypothesized that adiponectin may function as a lipid binding protein. Indeed, we found that recombinant adiponectin bound to various anionic phospholipids and sphingolipids, including phosphatidylserine, ceramide-1-phosphate, and sulfatide. The globular head-domain of adiponectin was necessary and sufficient for lipid binding. Adiponectin oligomerization was also observed to be critical for efficient lipid binding. In addition to lipids in liposomes, adiponectin bound LDL in an oligomerization-dependent manner. Other C1qTNF-related protein (CTRP) family members Cbln1, CTRP1, CTRP5, and CTRP13 also bound similar target lipids in liposomes. These findings suggest that adiponectin and other CTRP family members may not only function as classical hormones, but also as lipid binding opsonins or carrier proteins.

## INTRODUCTION

Since the discovery of adiponectin as a metabolic hormone secreted by adipocytes, there have been many studies describing the protective effects of adiponectin against cardiovascular disease, insulin resistance, and inflammation in sepsis and metabolic disease (1–8). Many of these discoveries have been complemented by studies on the adiponectin receptors Adipor1 and Adipor2 (1, 9), which are thought to mediate most of the myriad effects of adiponectin by inducing AMPK signaling, PPARα transcriptional programs, and receptor-intrinsic ceramidase activity (10). The ability of adiponectin to reduce tissue ceramide levels has been posited to explain its insulin sensitizing and anti-inflammatory properties (11–13). Adiponectin also has been shown to bind a separate GPI-linked receptor T-cadherin (14), which has recently been shown to also reduce tissue ceramide levels by triggering the release of exosomes (15). These findings together suggest a role for adiponectin in ameliorating metabolic disease by altering tissue lipids, especially ceramides, via receptor binding and activation of downstream intracellular processes.

However, several features of adiponectin biology are not readily consistent with the prevailing view that adiponectin functions only as a classical hormone. First, the concentration of adiponectin in serum is in micromolar range (16), which is in the range of carrier proteins such as IGF-binding proteins and retinol binding proteins; most other hormones, including insulin and leptin, circulate in the nanomolar range. Another unexplained feature of adiponectin is the presence of multimers and cleaved forms, which appear to have different physiological effects. Adiponectin spontaneously forms trimers, which assemble into higher-order dimers of trimers, and 4-5-mers of trimers. These are, respectively, referred to as low molecular weight (LMW), medium molecular weight (MMW), and high molecular weight (HMW) complexes.

Adiponectin has also been shown to be cleaved in serum by elastases and thrombin at the juncture between the globular C1q-like head-domain and the collagenous tail-domain. Interestingly, the globular head-domain appears to bind AdipoR1/2 and activate AMPK more strongly than full-length protein (17, 18); however, the quantities of HMW multimers are more strongly associated with insulin sensitivity (19, 20), have a longer half-life in circulation (16), and are selectively bound by T-cadherin over globular/trimeric forms (14). The reasons for these differences are not well-explained by the current paradigm of a purely endocrine function of adiponectin. Moreover, several studies have also shown an important protective effect of adiponectin in clearing apoptotic cells and preventing development of autoimmunity (21, 22). This effect, thought to be mediated in part via binding to calreticulin, appears to lie completely outside the metabolic realm of adiponectin biology, and a clear relationship between these processes has been lacking.

While the structural similarity of adiponectin and C1q has been previously noted (23), the possibility that these proteins may have similar functions has not been explored. C1q is best known for its role in innate immunity as an opsonin and the initiating protein in the classical complement cascade. It binds a wide variety of proteins, in particular the F_C_ domain of various immunoglobulins, C reactive protein, and β-amyloid, as well as carbohydrates and anionic phospholipids, including lipopolysaccharide, cardiolipin, and phosphatidylserine (23). Unlike adiponectin, which forms homotrimers, C1q is made of a heterotrimer of three distinct subunits (C1qA, B, and C), which have slightly different affinities to the various described ligands. The overall structure of adiponectin, C1q, and other C1q-TNF related protein (CTRP) family members, however, are strikingly similar in the C-terminal globular head-domains, as revealed by domain analysis and crystallographic studies (24). CTRP family proteins also bear resemblance to collectins, including surfactant proteins and lectins, and ficolins. Of note, these are also known to bind a range of phospholipids and carbohydrates (25–27).

Given these structural similarities, the involvement of adiponectin in apoptotic cell clearance, its unexplained high circulating levels, its relation with tissue ceramides, and the known effects of ectopic lipid deposition on insulin resistance and metabolic disease (28, 29), we hypothesized that adiponectin may bind lipids directly, serving as a physiological lipid opsonin. Using different biochemical approaches, here we find that adiponectin is a lipid binding protein with broad specificity to anionic phospholipids and sphingolipids, including phosphatidylserine, sulfatide, and ceramide-1-phosphate. These findings suggest a modified model of adiponectin function as a lipid-based opsonin that may promote lipid clearance through the ceramidase activity of the AdipoR1/2 receptors.

## RESULTS

### Adiponectin binds a variety of anionic phospholipids and sphingolipids

For initial screens, recombinant murine adiponectin with a V5-6xHis C-terminal tag was cloned and expressed in Expi293 cells, harvested in cell supernatant, and tested for binding to lipids on nitrocellulose strips either prepared in-house or obtained commercially (Echelon, Avanti). By this method, adiponectin was found to bind a variety of anionic phospholipids and sphingolipids, including phosphatidic acid (PA), phosphatidylserine (PS), phosphatidylinositol-phosphate (PIP), phosphatidylinositol-4,5-bisphosphate (PIP_2_), phosphatidylinositol-(3,4,5)-trisphosphate (PIP_3_), cardiolipin (CL), sulfatide (ST), ceramide-1-phosphate (Cer1P), and dihydroceramide-1-phosphate (DHCer1P) (Figure 1A). Of these, PS, CL, ST, and Cer1P were of interest given that they can be exposed in various conditions on the outer leaflet of cells, where they might interact with adiponectin physiologically. The binding to Cer1P was especially interesting given the relationship of adiponectin with ceramide metabolism (10, 11). Notably, there was little or no binding to triglycerides, diacylglycerol, cholesterol, neutral sphingolipids and phospholipids— including ceramide and phosphatidylcholine—or single-tailed sphingosine derivatives.

**Figure 1.**
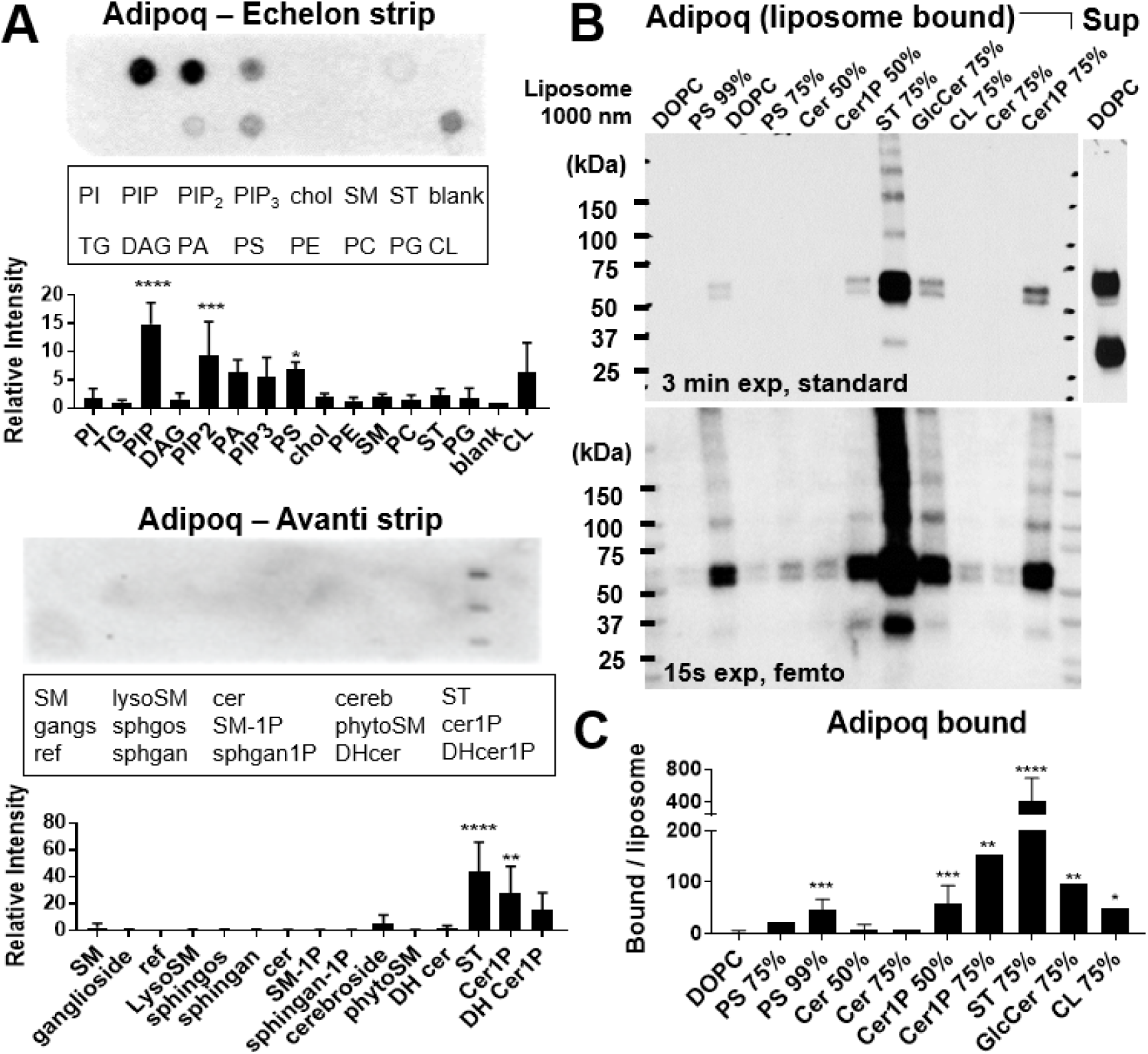
Adiponectin binds multiple negatively charged phospholipids and sphingolipids. (A) Binding of recombinant murine adiponectin to lipids on lipid strips, quantified at right. Density normalized to blank or reference spot. n ≥ 3 experiments. (B) Binding of adiponectin to liposome lipids in pulldown assay. Adiponectin in pellet fractions (liposome bound), and excess in supernatant (Sup) shown. Percentage of target lipids in liposomes indicated above each lane. Two exposures shown to highlight strong and weak bands. (C) Densiometric quantification of adiponectin V5 signal in liposome pulldown Western blots over multiple experiments. Band densities normalized to liposome rhodamine fluorescence and averaged over n ≥ 5 experiments. Error bars represent S.D. from the mean. Statistical significance of binding over blank/reference (A) or over DOPC (C) assessed by one-way ANOVA with Dunnett’s correction. * p < 0.05, ** p < 0.01, *** p < 0.001, **** p < 0.0001.

To rule out a role for other proteins in the transfected cell supernatant in bridging binding between adiponectin and lipids, lipid binding on nitrocellulose was assessed after isolation of recombinant tagged adiponectin by Ni NTA affinity chromatography (Supplementary Figure S1). Both purified and unpurified recombinant adiponectin showed comparable results, suggesting a direct interaction. Notably, deletion of the C1q domain also was able to abrogate binding, ruling out non-specific binding secondary to purification conditions. Purified recombinant protein was then used for all subsequent experiments.

Because lipid strip assays may show non-physiological interactions, given the random orientation of the dried lipids, binding of adiponectin to a subset of lipid targets was validated using liposomes, in which lipids are presented to proteins in a more physiological orientation and context, using a liposome pulldown assay. Briefly, lipids of interest incorporated into rhodamine-labeled, sucrose-loaded 1000 nm dioleylphosphatidylcholine (DOPC) liposomes were incubated with purified recombinant protein, washed, collected by density centrifugation, and analyzed for bound protein by Western blot. The protein signal was normalized to liposome rhodamine fluorescence, and unbound protein in the supernatant was monitored for comparison. By this method, adiponectin was again seen to bind various anionic phospholipids and sphingolipids (Figure 1B). Significant binding was observed to PS, Cer1P, glucosylceramide (GlcCer), and CL, but binding was remarkably strong to ST, largely corroborating the lipid strip data, but also revealing interesting differences. Of note, in all pulldown conditions, a large excess of recombinant protein remained unbound in the supernatant, irrespective of liposome type, despite differences in protein binding (Supplementary Figure S2).

In addition, we observed that selective binding of adiponectin to target lipids in liposomes required a high concentration of target lipids, >50% for PS and Cer1P in liposomes. Binding to ST, though stronger, still required >10% lipids for efficient binding (Supplementary Figure S2). Dot blot data also suggested a dependence of adiponectin binding on tail length of the lipid ligand (Supplementary Figure S3A). Interestingly, despite previous literature noting coordination of Ca^2+^ ions by adiponectin, lipid binding to strips or liposomes was not affected by the presence of calcium in the binding buffer (Supplementary Figure S3B-C).

### Lipid binding occurs via the C1q domain of adiponectin, and is affected by mutation of R134

Loss of adiponectin lipid binding of upon deletion of the C1q domain suggested that this domain may be important for interactions with lipids. To further characterize the necessity and sufficiency of the C1q domain for lipid binding, “headless” C1q-domain deleted mutant (C1qΔ) adiponectin (residues 1-110), and “tailless” collagenous-domain deleted mutant (collΔ) adiponectin (residues 1-44, 111-247) were tested for binding to lipids in lipid strips (Supplemental Figure S1) and liposomes (Figure 2A). By both methods, the C1q head domain was observed to be necessary and sufficient for lipid binding to PS, ST, and Cer1P targets. With liposomes, however, a clear defect in lipid binding was observed in the collΔ mutant compared to WT adiponectin, suggesting the head domain alone binds lipids with less avidity than the full-length protein.

**Figure 2.**
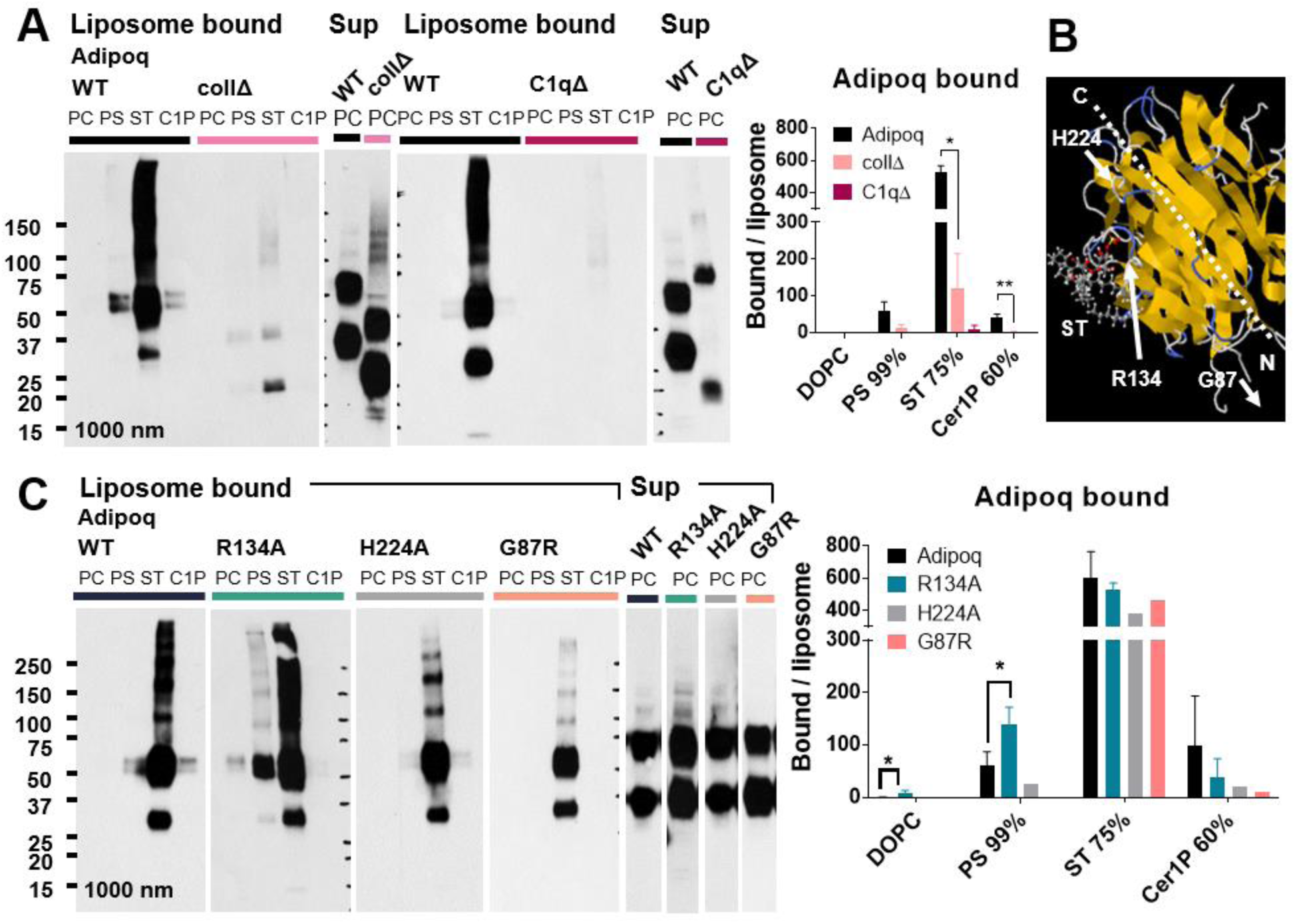
Lipid binding properties of adiponectin are mediated by the C1q domain, and are altered by R134A mutation. (A) Binding of WT adiponectin, and collΔ, C1qΔ mutants to liposome lipids by pulldown assay. Liposome composition labeled above each lane: PC = 99% DOPC, PS = 99% PS, ST = 75% ST, C1P = 60% Cer1P. Liposome-bound and excess unbound protein (Sup) shown for comparison. Normalized band density averaged over n ≥ 2 experiments graphed at right. (B) Position of adiponectin residues selected for site-directed mutagenesis relative to a potential ST binding site predicted using SwissDock on a crystal structure of a single-chain trimer of globular recombinant human adiponectin (4DOU) (31, 32). (C) Lipid binding of WT adiponectin and point mutants by liposome pulldown. Quantification as above over n ≥ 2 experiments. Error bars represent S.D. from the mean. ** p < 0.01, * p < 0.05.

Given the involvement of the C1q domain in lipid binding and the preference of adiponectin for phospholipids and sphingolipids with anionic headgroups, we hypothesized that positive residues in the adiponectin C1q head domain may be involved in ligand selectivity. Previous work characterizing the binding of PS to C1q suggested the R111 residue of human C1qC (R139 in the consensus sequence) may interact with the polar serine headgroup (30). Sequence alignment analysis revealed an analogous arginine (R134) in murine adiponectin conserved in human adiponectin. Docking studies of ST on the adiponectin C1q head domain (4DOU) (31) using SwissDock (32) also showed a conformation with estimated ΔG = −10.24 kcal/mol where the negative sulfate group lies in close proximity to the R134 residue (Figure 2B). Docking of Cer1P showed a similar conformation with an estimated ΔG = −8.75 kcal/mol.

To test the effect of R134 on selectivity for anionic lipid ligands, we introduced an R134A mutation in the WT protein, as well as an H224A mutation targeting the top rather than the side of the C1q domain, and a G87R mutation, which lies outside the C1q domain but has a weak association with diabetes in humans (19). Interestingly, while the R134A mutation did not abrogate lipid binding to anionic lipid ligands, it appeared to increase the relative affinity of adiponectin to PS and decrease binding to Cer1P (Figure 2C). The H224A mutation had essentially no effect, suggesting that this residue does not contribute significantly to lipid binding. The effect of the G87R mutation was subtle, but marginally decreased binding of both PS and Cer1P, while sparing binding to ST.

### Oligomerization of adiponectin is required for lipid binding

Upon inspection of the adiponectin bands in the pulldown assays, we noted a predominance of the upper 60 kDa band in liposome-bound protein versus an almost equivalent if not flipped distribution of the 30 kDa and 60 kDa bands in the unbound supernatant (Figure 3A). As adiponectin monomers run around 30 kDa, the 60 kDa band, which collapses to 30 kDa in reducing conditions, was thought to correspond to a disulfide-bonded adiponectin dimer. Since adiponectin trimers are held together primarily by hydrophobic interactions (33), we surmised that the apparent dimer band corresponds to inter-trimer dimers linked at the N-terminal cysteine, like those seen in C1q (34), in HMW and MMW adiponectin oligomers (Figure 3A). The predominance of the dimer band in liposome-bound fractions thus was hypothesized to reflect preferential lipid binding by higher molecular weight adiponectin oligomers. This, along with the differences in lipid binding between full-length and collΔ adiponectin, and in the physiological functions of various adiponectin oligomers, prompted an evaluation of the contribution of oligomerization to lipid binding.

**Figure 3.**
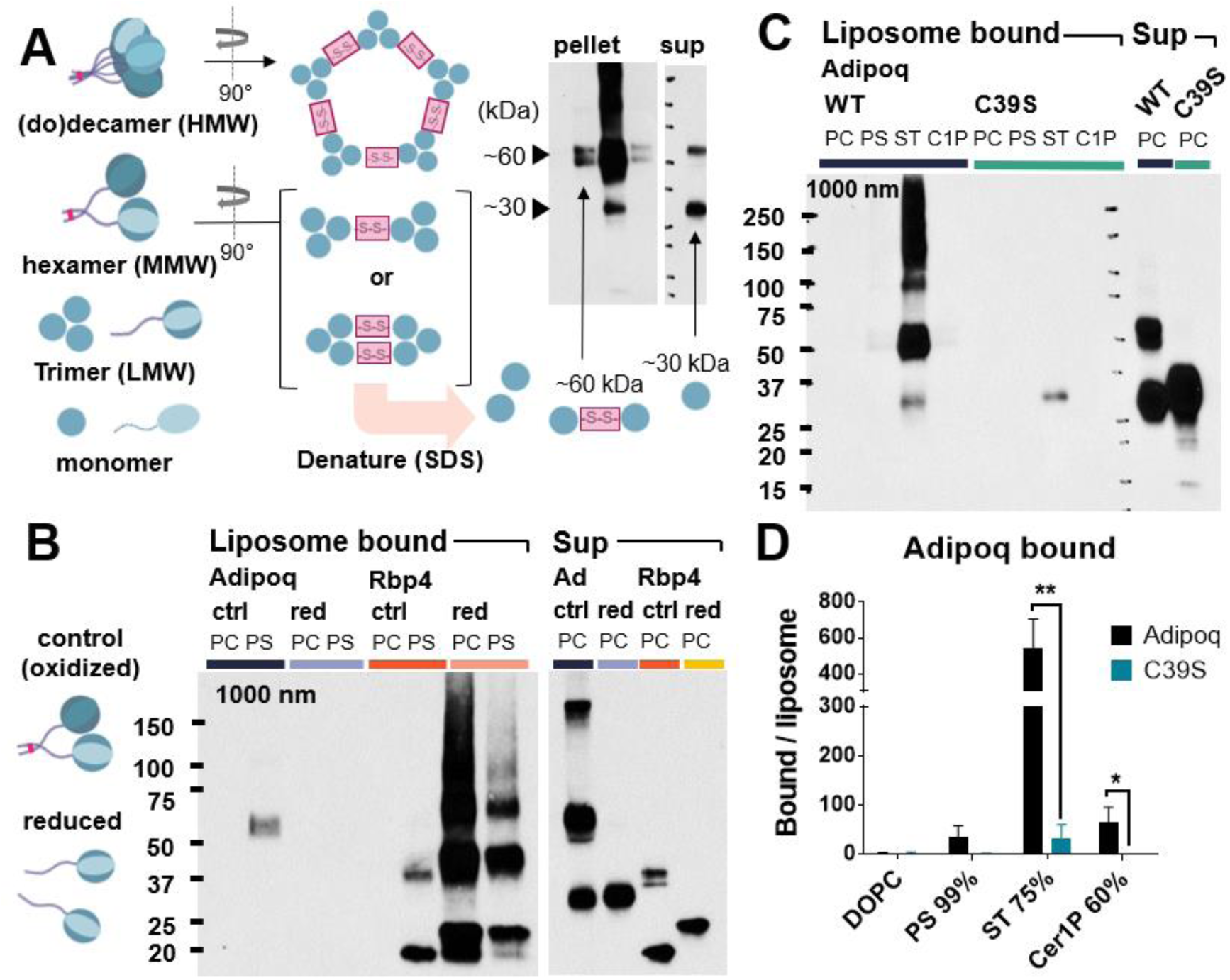
Oligomerization of adiponectin via disulfide bonds required for lipid binding. (A) Possible inter-trimer disulfide bond orientations in adiponectin multimers giving rise to observed ∼60 kDa and ∼30 kDa bands on non-reducing denaturing SDS-PAGE Western blot, which disrupts intra-trimer hydrophobic interactions. (B) Liposome binding of adiponectin and RBP4 in normal (oxidized) or reducing conditions with 2% β-mercaptoethanol. (C) Binding of WT and C39S adiponectin to lipids in liposomes. Excess unbound protein in supernatant shown at right. Quantification normalized to total liposome pellet fluorescence below, expressed as mean and S.D. over n = 3 experiments. ** p < 0.01, * p < 0.05. PC = 99% DOPC, PS = 99% PS, ST = 75% ST, C1P = 60% Cer1P.

Initial gel filtration analysis of purified recombinant murine adiponectin expressed from Expi293 cells showed three peaks consistent with the presence of HMW, MMW, and LMW adiponectin (Supplementary Figure S4). Having established the presence of higher molecular weight oligomers in isolated protein, we then tested whether reduction of disulfide bonds by β-mercaptoethanol would affect binding to target lipids in liposomes. Retinol binding protein 4 (RBP4), which was observed to have some binding to PS, was used as a control for potential non-specific effects of reducing conditions on lipid binding. Consistent with our hypothesis, binding of adiponectin to PS liposomes was completely abrogated by reduction of adiponectin (Figure 3B). Reduction also, as expected, eliminated the 60 kDa and other higher MW adiponectin bands. RBP4, which also had some binding to PS in the native state, conversely showed *increased* binding to PC and PS liposomes in the reduced state, suggesting that reducing conditions do not indiscriminately reduce lipid binding. Nonetheless, to further confirm the effect of adiponectin oligomerization on lipid binding without using reducing agents, site-directed mutagenesis was used to introduce a C39S mutation, which has previously been shown to abrogate adiponectin oligomerization (18). This mutation resulted in loss of the 60 kDa band even in non-reducing conditions, confirming that the 60 kDa band arises from oligomerization, and diminished binding of adiponectin to target liposomes (Figure 3C-D), corroborating the results with reduced and native protein. Of note, light binding is seen to liposomes with the strongest ligand, ST, indicating that binding is not completely abrogated by the mutation, but much reduced.

### Adiponectin binds lipids in physiological membranes and LDL

Our experiments thus far have investigated binding of adiponectin to selected target lipids at high concentrations in liposomes. While these studies serve to demonstrate that adiponectin binds lipids in a physically physiological setting, we wondered whether adiponectin binds lipids in more biologically relevant substrates in which the composition of lipids is more complex and the abundance of targets likely not as high as in synthesized liposomes, and what might be “natural” ligands of adiponectin in the body.

Since its discovery, adiponectin has also been associated with cardiovascular protection (3). Okamoto et al. first observed binding of adiponectin to areas of vascular injury (35), and adiponectin was later found to bind atherosclerotic lesions in mice (36, 37). Given these observations and the apparent lipid-binding property of adiponectin, we also tested whether adiponectin binds to low density lipoprotein (LDL) particles. To do this, native WT and C39S-mutated adiponectin were incubated with LDL in the presence of fatty-acid free BSA block, then subjected to flotation through an iodixanol gradient. The amount of LDL-associated adiponectin dragged upward through the gradient was then assessed by Western blot and aligned with Sudan black signal marking the LDL layer. As shown in Figure 4A, a substantial amount of WT adiponectin was indeed observed in the middle fractions with high Sudan black signal, suggesting binding to LDL. A large proportion, however, remained unbound in the lower fractions, as was also seen in liposome pulldown assays. Notably, flotation of adiponectin was lost with the C39S mutant, suggesting that adiponectin binding to LDL is via LDL lipids. The same pattern was observed with native and reduced WT protein (Supplementary Figure S5).

**Figure 4.**
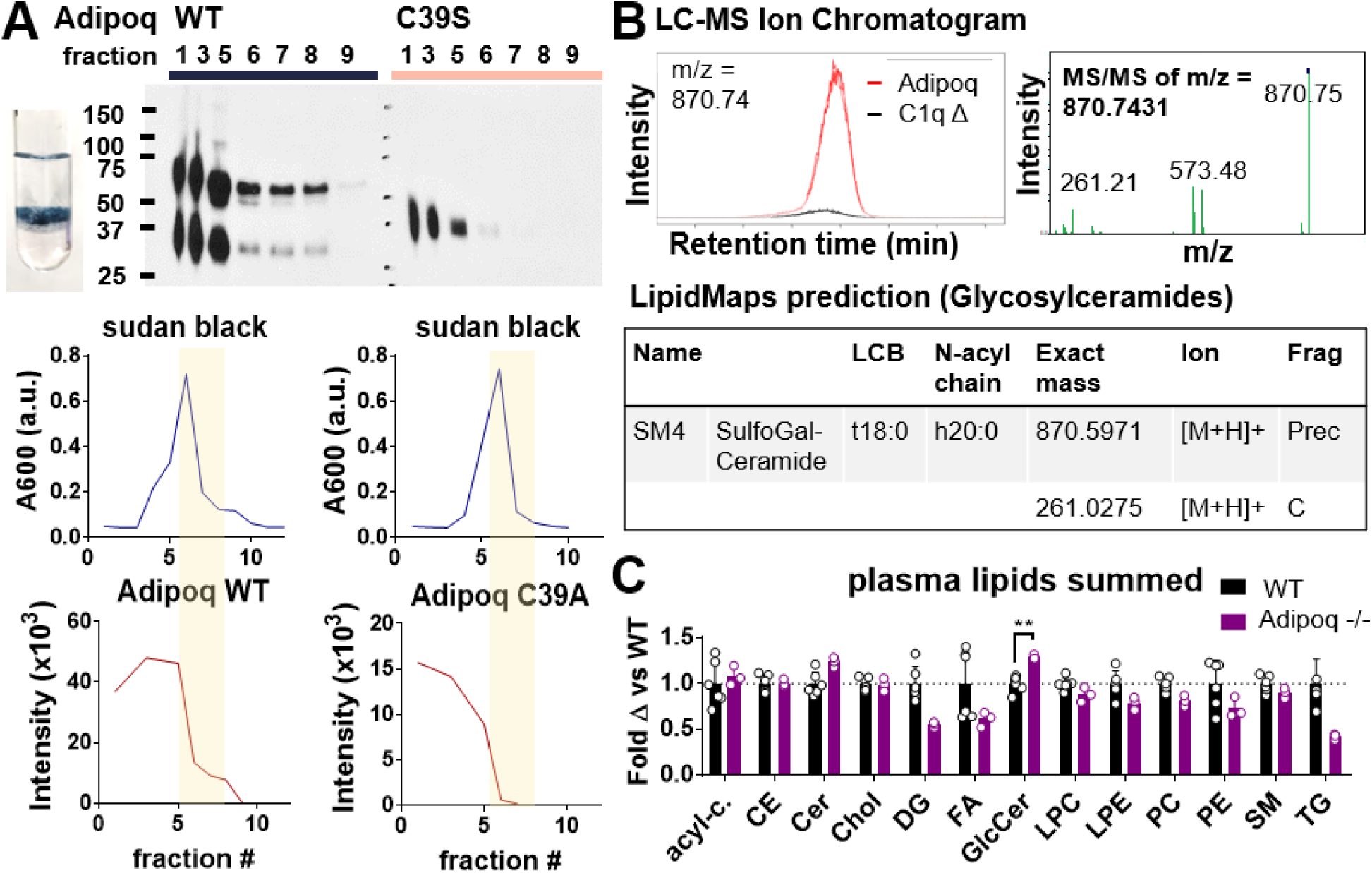
Adiponectin binds lipids in physiological context. (A) Adiponectin binds LDL lipids in oligomerization-dependent manner. (Top) Adiponectin protein in sequentially eluted iodixanol gradient fractions in LDL flotation assay, as detected by Western blot. Fraction number above each lane. Inset photograph shows banding pattern of LDL, stained with Sudan black, in the iodixanol gradient after ultracentrifugation. (Bottom) Quantification of Sudan black absorbance (upper) and adiponectin protein in Western blots (lower) in sequential gradient fractions. Yellow bars highlight Sudan black peaks corresponding to the location of LDL in the gradient. Representative blots and graphs of n = 3 experiments shown. (B) Adiponectin isolated from cell culture associated with sulfatide. (Left) LC-MS lipid peak with high binding to adiponectin but no binding to C1qΔ deletion mutant or Rbp4 control protein. (Right) MS/MS fragmentation of the m/z = 870.74 peak. (Bottom) Characteristics of predicted lipid with the observed fragmentation pattern found using the LipidMaps glycosylceramide search tool. (C) Shotgun lipidomics analysis of plasma of WT and Adipoq KO mice. Peak intensity reads, normalized to that of the corresponding internal standard, summed over all identified lipids within a category and graphed as fold change relative to the WT average for that lipid. Error bars reflect S.D. from the mean. Open circles represent values from individual mice. ** p < 0.01, * p < 0.05. Unmarked bars were found to be not significantly different from WT.

We also wondered whether adiponectin may bind target lipids in cell membranes. To assess this, we performed an untargeted LC-MS screen to characterize what lipids, if any, are associated with adiponectin simply upon isolation from transfected Expi293 cell supernatant. As there are no serum or protein additives in Expi293 media, and expressed adiponectin is in contact with cells for days until isolation, any bound lipids presumably are derived from the surrounding cells, potentially via extraction from plasma membranes, or released in exosomes or other membranous debris. Of note, non-specific adsorption of lipids to proteins is frequent and leads to a high rate of false positives by this method. To circumvent this, we performed LC-MS with three protein preparations: WT adiponectin, the C1qΔ mutant, and RBP4. Only significant peaks specific to WT adiponectin in both WT to C1qΔ and WT to RBP4 comparisons were considered for further testing; common peaks were considered non-specific and ignored. A number of WT adiponectin-specific peaks fit this criterion (Supplementary Figure S6), 11 of which were enriched at least 2.5-fold in WT adiponectin in both comparisons, had p values < 0.01, and peak max intensity ≥ 20,000. Many of these could not be mapped to a known compound using the XCMS-MRM platform algorithms (38). Of the remaining, two were mapped to triglyceride or phosphatidylcholine, one to PIP_2_, one to a glucosaminyldiphosphoenol species, and one to galactosylceramide or sphingomyelin. This last peak (m/z = 870.7) was the most interesting to us given our previous lipid binding data, and was selected for further ms/ms fragmentation (Figure 4B). Fragment peaks were searched against the METLIN ms/ms database and various LipidMaps libraries (39, 40). Of these, only the LipidMaps glycosylceramide prediction tool was able to map two of the ms/ms peaks to a single hit, matching the m/z = 870.7 and m/z = 261.2 ion peaks to the precursor and C-fragment ions (glycosidic bond fragment) of sulfogalactosyl-ceramide, also known as ST, corroborating the liposome lipid binding data, and demonstrating that adiponectin can bind ST in cell membranes.

Given these indications that adiponectin may bind physiologically relevant lipids, and our hypothesis that adiponectin acts as an endogenous lipid opsonin, we investigated whether any changes could be observed in lipid levels *in vivo*, especially in the plasma where adiponectin is found in high concentrations. To do this, plasma of WT and Adipoq KO mice were collected and sent for shotgun lipidomics analysis. Most detected lipids were not adiponectin ligands and were not significantly different between WT and KO mice; however, glucosylceramide species, which by liposome assay did bind adiponectin, were significantly elevated in KO mouse plasma (Figure 4C, Supplementary Figure S7). This suggests adiponectin lipid binding *in vivo* may play a role in reducing the levels of this lipid.

### Adiponectin does not specifically label dead cells

Given the role of adiponectin in clearance of apoptotic cells, and our observation of adiponectin binding to PS in liposomes, we investigated whether adiponectin might also bind PS in necrotic or apoptotic cells using flow cytometry. For this, purified preparations of WT and C39S adiponectin were used to stain a mixed-viability population of Expi293 cells co-stained with Annexin V and propidium iodide (PI). While recombinant WT adiponectin was able to clearly bind to dead (Annexin V+, PI+) cells and a fraction of apoptotic cells (Annexin V+, PI-), the same binding pattern was observed with C39S adiponectin, which was seen earlier to lack binding to PS (Supplementary Figure S8A-B). This suggests that adiponectin binds ligands exposed on dead cells other than PS, like calreticulin, or alternatively, that adiponectin recognition of PS on apoptotic cells may not require oligomerization.

### Adiponectin does not have intermembrane lipid transfer activity

Given the ability of adiponectin to bind lipids, we wondered whether adiponectin may function primarily by binding, like an opsonin, or have additional lipid extraction or transfer activity. While adiponectin lacks an obvious hydrophobic pocket that might mediate lipid transfer, the possibility was formally assessed using a loss-of-FRET quenching lipid transfer assay, where transfer of labeled lipid ligands from quenched donor liposomes to unlabeled acceptor liposomes is monitored by the appearance fluorescence over time. Experiments were performed with Cer1P, PS, and ST-containing donor liposomes, as well as ceramide-containing donor liposomes as a negative control. Activities of WT adiponectin, the C1qΔ mutant, collΔ mutant, and C39S mutant were also compared. Controls with protein with donor liposome alone, or donor and acceptor liposome without protein, were included in some cases to assess lipid extraction by protein without donation, or spontaneously lipid transfer, respectively. From these experiments, adiponectin was not found to have any lipid transfer activity for ceramide or Cer1P, PS (Supplementary Figure S9). Adiponectin also did not show lipid extraction or transfer activity for ST, although there was a high degree of spontaneous lipid transfer between donor and acceptor liposomes that confounded the assessment of transfer activity.

### CTRP family members bind a similar set of anionic phospholipids and sphingolipids

Adiponectin and C1q belong to the larger C1q-TNF related protein (CTRP) family, which consists of the C1q, emilin, C1q-like, and cerebellin (Cbln) subfamilies (41, 42). Various crystallographic studies have shown that CTRP family members share similar structures to adiponectin and C1q, forming trimers and oligomers of trimers (24), primarily differing in the length and complexity of the collagenous tail. The C1q regions are generally conserved, with relatively strong conservation of the hydrophobic structural elements of the β-strand jelly-roll, but more variability in the solvent exposed loops and superficial electrostatic patterns (24, 41, 42).

Having observed the ability of adiponectin to bind to various anionic phospholipids and sphingolipids, we wondered to what degree this property might be shared among CTRP family members, and whether their affinities for various lipids may differ. To test this, V5-6xHis tagged recombinant Cbln1(precerebellin 1), CTRP1 (GIP), CTRP5 (Myonectin), and CTRP13 (C1ql3), representing three of the four CTRP subfamilies (Figure 5A), were cloned, expressed, and isolated by Ni NTA affinity chromatography as for adiponectin (Supplementary Figure S8) and assessed for lipid binding by liposome pulldown. CTRP5 is a protein expressed in the retinal pigment epithelium and certain brain regions that belongs to the C1q subfamily with adiponectin and C1qA, B, and C (43). Cbln1 and CTRP1 are cerebellin subfamily proteins; Cbln1 is expressed in cerebellum and known to be involved in synaptic organization (44, 45), while CTRP1 is expressed peripherally in multiple tissues, including adipose, and was found to have a role in glucose and lipid homeostasis (46). CTRP13 is a member of the C1q-like subfamily expressed in brain cortical regions and retina with some role in regulating synaptogenesis (47). Interestingly, all the CTRPs cloned showed some degree of binding to PS, Cer1P, and especially ST (Figure 5B). As with adiponectin, this lipid binding appeared to depend on oligomerization mediated by disulfide bonds, as illustrated by loss of binding of reduced versus native CTRP1 (Figure 5C). Affinities to various lipids generally appeared to follow the pattern of adiponectin, but some variability was observed among family members. CTRP1 in particular had much stronger binding to PS relative to ST than any of the other proteins. Cbln1 also appeared to have a higher affinity for ST, as suggested by the high degree of binding to ST liposomes despite lower concentrations in the supernatant. In other blots, binding of Cbln1 to Cer1P and PS was similar to that of adiponectin (Figure 5D). Of note, the CTRPs cloned differed quite dramatically in their level of expression in Expi293 cells: CTRP5 and CTRP13 were poorly expressed relative to Cbln1, CTRP1, and adiponectin, which limited assessment of lipid binding.

**Figure 5.**
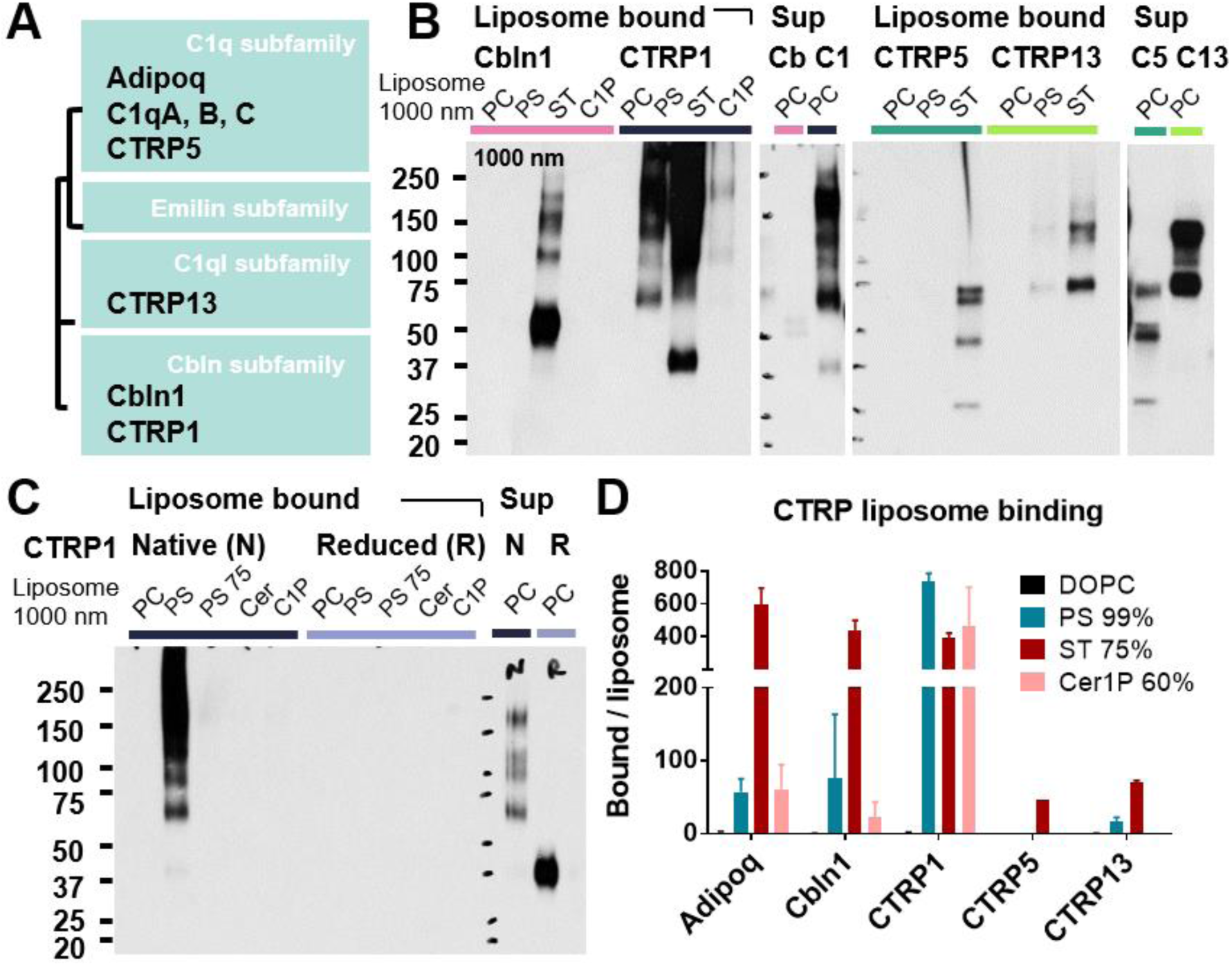
Other C1qTNF-related proteins also bind lipids in liposomes with similar but slightly different patterns of affinity. (A) Simplified tree diagram showing relationships between CTRP family members cloned for this study. (B) Liposome pulldown assays showing binding of Cbln1, CTRP1, CTRP5, and CTRP13 to liposomes with various target lipid ligands. Composition of liposomes indicated above. PC = 99% DOPC, PS = 99% PS, PS 75 = 75% PS, ST = 75% ST, C1P = 60% Cer1P, Cer = 60% C16 ceramide. (C) Loss of lipid binding of CTRP1 to target lipids in liposomes in reducing conditions. (D) Quantification of CTRP lipid binding in liposome pulldown blots. Band density normalized to liposome pellet rhodamine fluorescence. n.t. = not tested. Bars represent mean and S.D. over n ≥ 2 experiments.

The similar patterns of affinity for PS, ST, and Cer1P ligands of CTRP1 and R134A adiponectin, versus that of WT adiponectin and the other CTRP family members cloned, prompted an examination of the amino acid sequences to determine whether the observed differences might be explained by variation in arginine or other basic residues. Alignment of consensus sequences of human and murine C1qC, human and murine adiponectin, and murine Cbln1, CTRP1, CTRP5, and CTRP13 was performed with Uniprot (Supplementary Figure S10) and basic structural features were annotated using Uniprot data, existing crystal structures, and predicted structures predicted by the I-TASSER threading algorithm (48). In this alignment, the R134 residue of murine adiponectin was conserved across human adiponectin as well as mouse and human C1qC, as discussed (green triangle). This arginine was also partially conserved in Cbln1, CTRP5, and CTRP13 (substituted with a lysine in CTRP13, shifted by a few residues in Cbln1 and CTRP5), but was lost in CTRP1, which has a valine in the equivalent position and no nearby positive residues. Notably, all family members including CTRP1 have positive residues further upstream in the C1 solvent-exposed loop. The sequence of the CTRP1 C1 solvent-exposed loop is thus electrostatically similar to the R134A adiponectin mutant. Interestingly, the sequence alignment also showed strong conservation of the C39 cysteine residue of murine adiponectin across all family members (blue triangle), which might be expected to have an analogous role in protein oligomerization via disulfide bonds.

## DISCUSSION

In this study, we have demonstrated that adiponectin is a lipid binding protein that selectively binds various anionic phospholipids and sphingolipids, including PS, Cer1P, and ST, both in the context of synthetic liposomes, biological cell membranes, and LDL. Using site directed mutagenesis, we further determined that lipid binding occurs via the C1q domain, which is necessary and sufficient for binding; but also requires protein oligomerization via disulfide bonds at the C39 residue in the collagenous domain. Furthermore, we observed that the properties of the amino acids in the C-terminal solvent-exposed loops modulate selectivity for various lipid ligands, as mutation of the R134 residue changed the relative affinity of adiponectin for Cer1P and PS. More generally, we found that other representative CTRP family members bind to similar lipid ligands in a similarly oligomerization-dependent manner, and observed that natural variability in the amino acid composition of the first C-terminal solvent-exposed loop corresponds to differences in binding affinity to PS vs. Cer1P. Other solvent-exposed loops are also likely to contribute; for instance the (Y/F)FGGWP sequence in the C3 loop of CTRP5 was noted in previous crystallographic studies to form a hydrophobic patch, which may help stabilize lipid ligand binding (49). Given their lower sequence conservation, in comparison to the C1q β-sheet jelly roll core, it is possible that variation in these solvent exposed loops may have allowed the affinity of CTRPs for various ligands to be tuned over evolutionary time.

The observed lipid-binding properties of adiponectin with preference for anionic phospholipids and sphingolipids is consistent with the observed binding of C1q to CL and PS (23, 30). Though C1q lipid binding was not directly tested in this study, it is likely given its known lipid tropism that adiponectin and C1q bind similar lipid ligands. The conservation of this property across CTRP family proteins also fits with the more general carbohydrate- and phospholipid-binding properties of the structurally similar collectins and ficolins, such as mannose-binding lectin, surfactant protein, and others. For C1q, collectins, and ficolins, collectively referred to as “defense collagens,” ligand binding has a well-known role in neutralizing and opsonizing microbes. The current data on adiponectin, demonstrating lipid binding without apparent lipid transfer activity, suggest an analogous role as a physiological opsonin for extracellular lipids like LDL and other membranous debris enriched in target anionic phospholipids and sphingolipids. While adiponectin-mediated lipid uptake was not directly shown in this study (given confounding C1q secretion by phagocytes), a role for adiponectin in opsonization is corroborated by the relatively stable binding to target liposomes during pulldown experiments, which withstands multiple washes; lack of lipid transfer activity; and theoretical orientation of bound adiponectin with C1q heads in toward target lipids and collagen tails outward, available for recognition by phagocyte collagen and scavenger receptors. It is also supported by published data that adiponectin and C1q promote engulfment of apoptotic cells by a shared pathway (22).

While opsonization of lipids followed by uptake by phagocytosis is likely a shared function of C1q, adiponectin, and CTRPs, as well as some collectins and ficolins, each of these proteins may also have specific functions depending on their additional binding partners. C1q, mannose-binding lectin, and ficolins, for instance, can fix complement via C1r/s or MASP serine proteases (50), while adiponectin uniquely engages the Adipor1/2 receptors and generally does not activate complement in the native circulating state (51, 52) (Schematic 1). These specific functions may be key in differentiating the biological roles of the CTRPs, especially since they bind similar lipid ligands. In the case of C1q and adiponectin, with their similarly high abundance and ubiquitous distribution, these functions may largely complement each other, but could also compete. Adiponectin binding may mask a fraction of substrates from C1q, which despite having an important anti-inflammatory role in clearance of cellular debris, still may generate inflammatory cleavage products if a critical amount of C3b is deposited (53). By engaging Adipor1/2 receptors, inducing Appl- and Cav3-mediated receptor endocytosis, and stimulating ceramidase activity, adiponectin may additionally promote digestion and recycling of bound lipid cargo, removing them from the extracellular environment. Though the hydrolytic activity of the Adipor1/2 receptors was characterized for ceramides, a similar reaction can theoretically act on analogous bonds in other phospholipids and sphingolipids as long as they can enter the active site, which may be impeded by the anionic head groups of adiponectin target lipids, but could occur if Adipor1/2 receptor activity is paired with nearby phosphatases or glycosidases, or if the head groups are cleaved by other enzymes upon endocytosis. This assumes that lipid binding does not interfere with binding to the receptor, which cannot currently be determined as the residues of adiponectin involved in interaction with Adipor1/2 have not been reported. It is conceivable, however, especially in an adiponectin oligomer, that subunits not involved in lipid binding may be available for receptor ligation.

The common chemical properties of the lipids bound by C1q and adiponectin is notable, and begs the question of whether this is a feature of material that can or needs to be opsonized. PS for instance is well-known to mark apoptotic cells, and CL is associated with necrotic cell death, as it is typically sequestered in the inner membrane of the mitochondria. Less is known about Cer1P, a phosphorylated ceramide only present in minute quantities in whole cells, and ST, a sphingolipid present in most cell membranes and enriched in myelin sheaths. Both however have been observed to be secreted by cells into culture media (54), with ST being preferentially secreted in extracellular vesicles from senescent fibroblasts (55). ST also induces inflammation in microglia (56), and predisposes to seizures if accumulated in neurons (57).

Notably, the lipid binding by adiponectin and CTRPs was seen to depend both on target lipid density and protein oligomerization via disulfide bonds at C39. One explanation for this is that the individual lipid-protein interactions are relatively weak, and multiple interactions, i.e. through the multiple head subunits of an adiponectin oligomer, are necessary for sufficient avidity for stable binding. In physiological contexts, where target lipid abundance may be much lower (around 3-10% for PS and ST, and <0.1% for Cer1P in some membranes (58, 59)), patches of high target-lipid density may be assembled by lipoproteins, membrane rafts or scaffold proteins. This requirement for high target-lipid density and protein oligomerization for binding may be significant in providing a basis for regulating the extent and location of lipid binding by CTRPs.

Thus, our working model is that adiponectin HMW/MMW oligomers preferentially bind to high-density patches of anionic phospholipids/sphingolipids in LDL, apoptotic cell membranes, or other vesicular/membranous debris; and subsequently mediate their delivery to cells with adiponectin receptors for recycling by the receptor intrinsic ceramidase activity, or phagocytes that can recognize the collagenous domain (Schematic 2). This removes those lipids from extracellular spaces in tissue or vessels where they can cause issues with insulin sensitivity, inflammation, and/or inappropriate complement fixation after deposition of C1q. LMW/globular adiponectin may then complement this opsonization activity by promoting receptor activation without lipid delivery, thus priming the metabolic state of the tissue for lipid oxidation. This model is corroborated by the lipidomics analysis of plasma from WT and Adipoq -/- mice. It is also consistent with the known metabolic effects of adiponectin and suggests a more complete understanding of how it may achieve these effects. For instance, the protective effect of adiponectin against insulin resistance may result from targeting of adiponectin-bound lipids to macrophages and AdipoR1/2-expressing cells and improving clearance of pathogenic ectopic lipid debris, in addition to simple receptor activation. Clearance of LDL facilitated by adiponectin binding may also explain how adiponectin slows development of atherosclerosis in addition to directly limiting inflammation in macrophages (3, 60). The findings have interesting implications for the role of adiponectin-mediated lipid binding in metabolic disease, given the currently unexplained association of HMW vs LMW adiponectin with protection against Type 2 diabetes and metabolic syndrome (19, 20), the observed induction of and binding to exosomes by adiponectin (15, 61), and the role of exosomes in adipose tissue biology (62). The generalizability of this model to other CTRPs and the role of lipid binding in their function also remains an open question. Continuing research in this area will be instrumental in building a clearer understanding of extracellular lipid handling and its role in organismal health and disease.

## MATERIALS AND METHODS

Additional methods and technical details are available in the Supplementary Information.

### Protein expression and isolation

cDNAs for murine adiponectin and CTRPs were derived from adipose, brain, and other cDNA libraries and cloned into pcDNA 3.1 vectors with V5 6x His tags at the C-terminus. Proteins were then expressed using the Expi293 mammalian cell expression system. On day 5 post transfection, protein was harvested, passed through a 0.22 μm-pore filter to remove residual cellular debris, then incubated with Ni NTA beads overnight. Beads were then washed and eluted using imidazole in HNC isolation buffer (25 mM HEPES, 150 mM NaCl, 1 mM CaCl_2_, pH 7.4). Protein fractions were collected and concentrated using Amicon-Ultra 3- or 10-kDa molecular weight cut-off centrifugal spin filters (Millipore), then washed three times with HNC buffer to remove imidazole. A280 of the final isolate was measured by nanodrop. Protein was then kept at 4°C until use or snap frozen in liquid nitrogen and kept at −80°C for longer storage.

### Dot blot

Dot blots were either obtained commercially from Echelon Biosciences and Avanti Polar Lipids, or made from candidate lipids (obtained from Avanti Polar Lipids) spotted on 0.45 μm-pore nitrocellulose strips (2 μg/spot); 2 μl of anti-V5 antibody (Invitrogen, 46-0705) sometimes spotted as positive control. Dot blots were first blocked with 3% fatty acid-free BSA in TBST, then incubated with tagged, expressed protein in unpurified cell culture supernatant overnight. Blots were subsequently washed and probed with rabbit anti-6x His primary antibody (1:1000), followed by goat anti-rabbit peroxidase-conjugated secondary antibody, then developed using chemiluminescence substrate.

### Liposome pulldown assay

In brief, 1000 nm sucrose-containing liposomes of varying lipid composition, with 1% rhodamine-PE for visualization, were synthesized (see Supplementary Information for extended methods). Liposomes (final concentration 0.5 mM) were then incubated with 10 μg/mL isolated recombinant protein in the presence of 10 mg/mL fatty acid-free BSA block in HNC buffer at 37°C for 1 hr. Reduced protein, when used, was prepared by adding β-mercaptoethanol to a final concentration of 2% to the 10 μg/mL recombinant protein stock before addition to lipids. After binding, liposomes were diluted with 500μl of HNC, centrifuged at 15000 rpm × 15 min, then washed with an additional 500 μl HNC. Finally, pellets were resuspended in 25 μl HNC and added to a black opaque low-adsorption 96-well assay plate (Corning 3991) containing 3 μl 10% Triton X-100, and rhodamine fluorescence was measured. Samples were then denatured with 5x SDS non-reducing sample buffer, boiled, and analyzed by Western blot for the V5 tag. For each protein in each experiment, liposome-bound samples were run alongside excess unbound protein in supernatant, diluted 1:20, for comparison.

### LDL flotation assay

As in liposome pulldown assays, human LDL (final concentration 1 mg/mL) were incubated with 10 μg/mL isolated recombinant protein in the presence of 10 mg/mL fatty acid-free BSA block and 1 mg/mL Sudan black in HNC buffer at 37°C for 1 hr. Reduced protein prepared as above. After incubation, 50 μL of the LDL and protein mixture was added to 280 μL OptiPrep iodixanol solution to make a 330 μL, 1.25 g/mL layer, which was overlaid with 330 μL 70% OptiPrep, 670μL 36% OptiPrep, and 670 μL HNC buffer. Pairs of protein gradients were then ultracentrifuged at 50000 rpm x 20 hrs at 4°C with no brake on an Optima TL benchtop ultracentrifuge (TLS-55 rotor). Fractions were collected by puncture from the bottom of the tube, and absorbance of fractions at 600 nm was measured. Selected fractions were then collected into 5x non-reducing SDS sample buffer and analyzed by Western blot for V5.

### Protein-bound lipid analysis by liquid chromatography-mass spectrometry (LC-MS) and tandem mass spectrometry (MS/MS)

Expressed proteins were concentrated to 80-100 ul using an Amicon-Ultra 3 kDa MWCO spin filter, then snap frozen in liquid nitrogen and stored at −80°C until shipping. Subsequent LC-MS and MS/MS analysis was performed by the Scripps Center for Metabolomics and Mass Spectrometry. Samples were extracted 50:50 methanol:acetonitrile (ACN) and lipid extracts were analyzed by LC-MS (Bruker Impact II Q-TOF coupled to an Agilent 1200 LC stack). Positive ions were analyzed using XCMS software. MS/MS analysis was performed on m/z = 870.74. The resulting MS/MS spectral peaks were then searched against various databases in METLIN and LipidMaps. The prediction for sulfatide was found using the glycerophospholipid search tool in LipidMaps (http://www.lipidmaps.org/tools/ms/glycosylcer_gen.html).

### Plasma collection and lipidomics

8-10-week old male WT C57B6/J (JAX Stock No. 000644) and Adipoq -/- mice (B6;129-Adipoqtm1Chan/J, JAX Stock No. 008195) were housed in an SPF-facility with regulated 12 hr light-dark cycles with *ad libitum* access to standard chow and water. Whole blood was harvested by retro-orbital bleeding, and plasma was isolated using lithium-heparin coated plasma separator tubes (BD 365985) and stored at −20°C. Lipidomics was performed by the West Coast Metabolomics Center at UC Davis. In brief, extraction of samples was performed in MTBE with addition of internal standards, followed by ultra-high-pressure liquid chromatography on a Waters CSH column, interfaced to a QTOF mass spectrometer. Data were collected in both positive and negative ion mode, and analyzed using MassHunter (Agilent).

### Quantification and Statistical Analysis

Films and images were quantified using FIJI software. For all liposome pulldown assays, band densities were normalized to total pellet rhodamine fluorescence measured before lysis. Averages, standard deviations (S.D.), and statistical significance were calculated in GraphPad Prism. Students t-tests (two-tailed) were used for single comparisons between groups. Where multiple comparisons were made to the same control group, ANOVA with Dunnett’s correction for multiple testing were used. p-values less than 0.05 were considered significant.

## AUTHOR CONTRIBUTIONS

RM conceived of the project along with JJY, who designed and executed all the experiments performed and wrote the manuscript. JL suggested the methods for the initial experiments, and XB contributed technical expertise instrumental in the studies of liposome binding.

## ACKNOWLEDGEMENTS

The authors would like to thank the members of the Medzhitov lab, De Camilli lab, Cresswell lab and the Yale Department of Immunobiology for contribution of ideas, comments, tools, and reagents. Special thanks to Dr. Pietro De Camilli for guidance and reagents, with support from NIH grant DA018343. JJY is supported by the Yale School of Medicine Medical Scientist Training Program training grant (GM007205). XB is supported by the Human Frontier Science Program Long-Term Fellowship. JL is supported by Jane Coffin Childs Fellowship. RM is an investigator of the Howard Hughes Medical Institute.

---

**Schematic 1.**
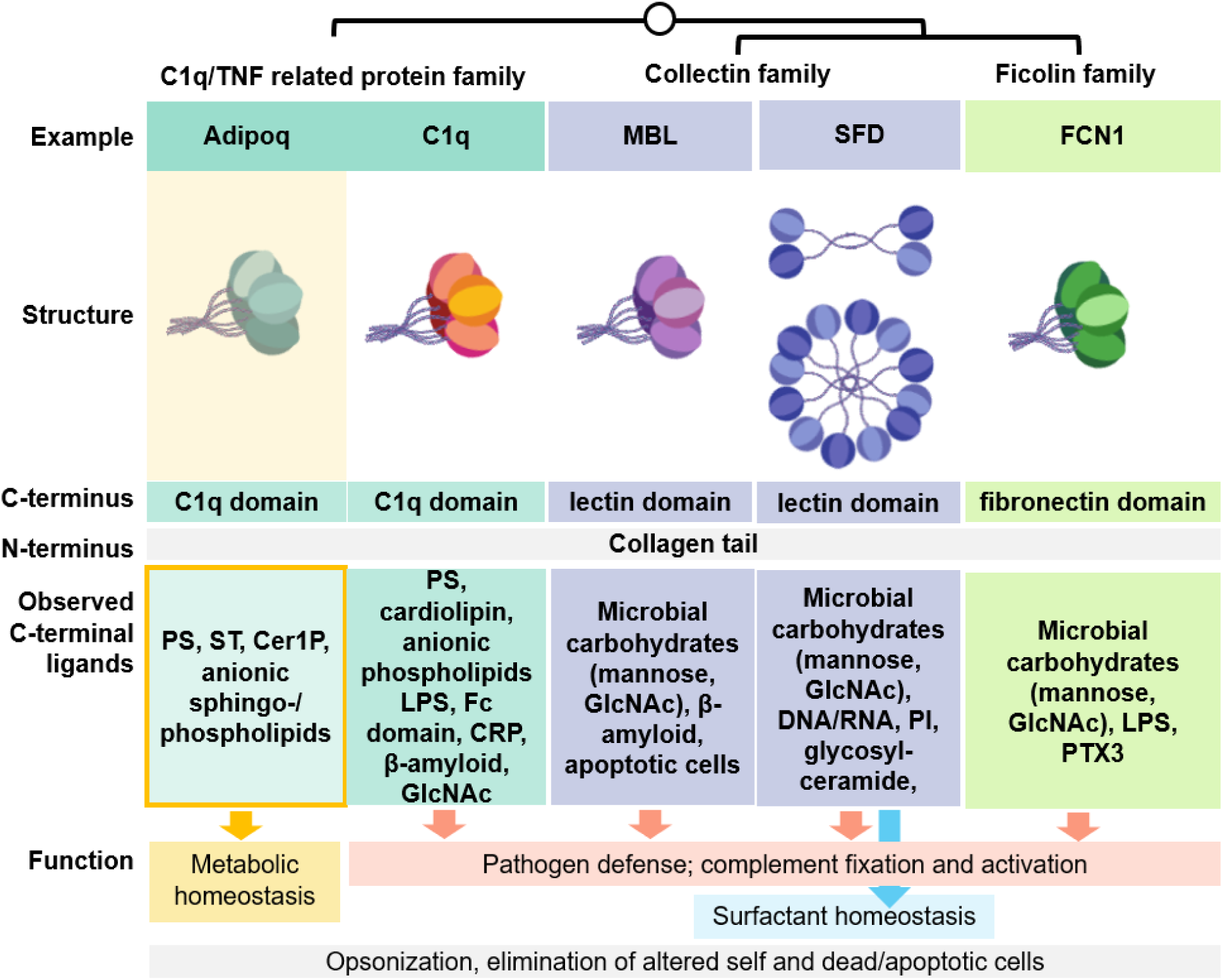
Binding of adiponectin to anionic sphingo/phospholipids fits with the known structural and ligand-binding features of C1q, collectins, and ficolins, but fills a unique functional niche in metabolism. Similarities and differences between various CTRP, collectin, and ficolin family proteins in terms of oligomeric structure, C- and N-terminal domains, C-terminal binding targets, and currently understood function. Possible shared features, like binding, opsonization, and elimination of self-targets indicated in grey. Additional possible unique outcomes for each protein indicated in color-coded boxes. Adipoq = adiponectin, MBL = mannose-binding lectin, SFD = surfactant protein D, and FCN1 = ficolin 1.

**Schematic 2.**
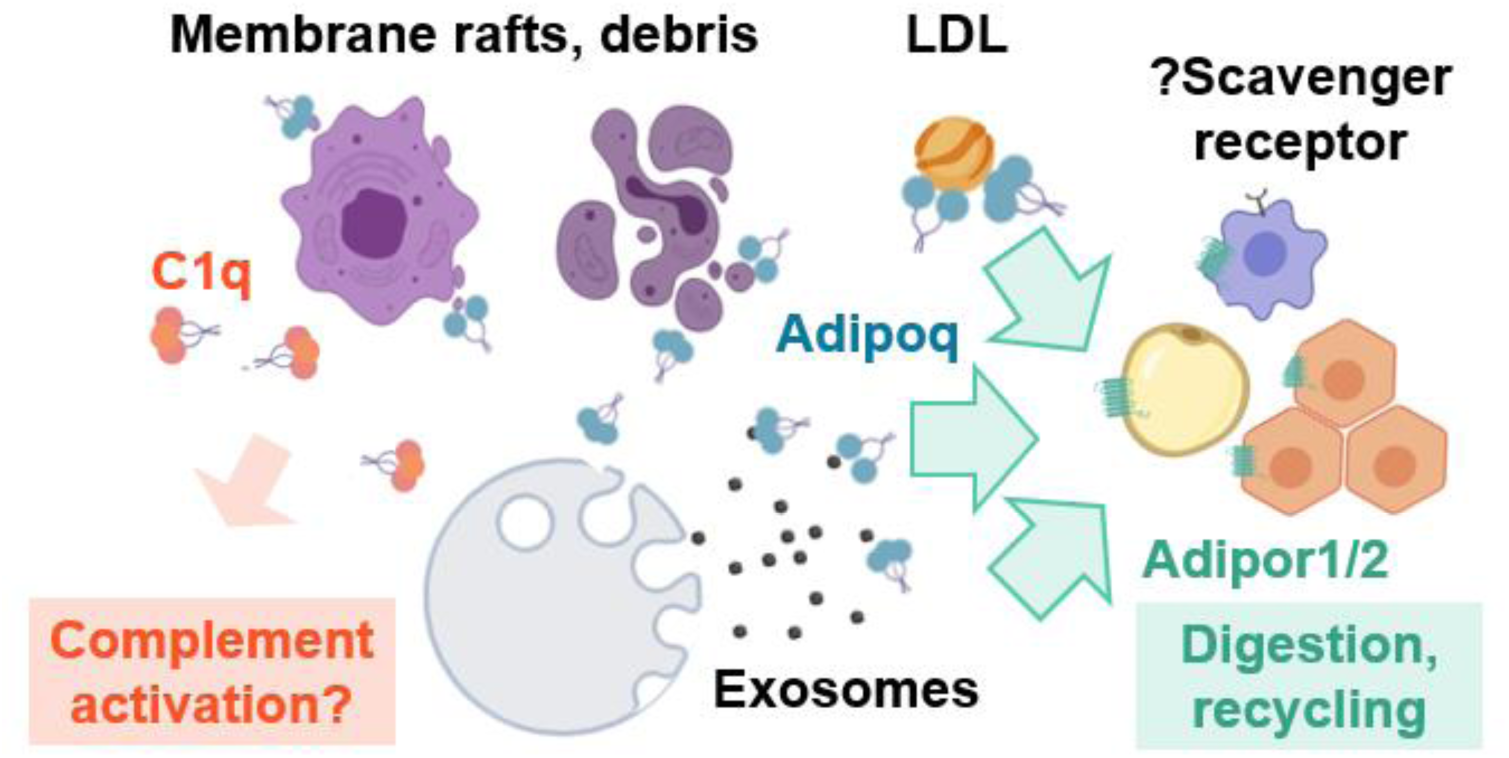
Binding of negatively-charged lipids by adiponectin may promote clearance and limit complement-mediated inflammation. Adiponectin, C1q, and other CTRP family proteins appear to bind similar target lipids, including PS, Cer1P, and ST potentially found in lipoproteins, exosomes, and membranous debris. Unlike C1q, however, adiponectin does not generally activate the complement cascade, and may mediate clearance by binding AdipoR1/2. The collagenous N-terminal domain may also permit uptake via macrophages scavenger receptors. In this way, adiponectin may both promote clearance and recycling of lipid debris, and dampen activation of complement by competing for overlapping lipid targets.

## REFERENCES

1. Kadowaki T, et al. (2006) Adiponectin and adiponectin receptors in insulin resistance, diabetes, and the metabolic syndrome. J Clin Invest 116(7):1784–1792.

2. Wang Z V., Scherer PE (2016) Adiponectin, the past two decades. J Mol Cell Biol 8(2):93–100.

3. Okamoto Y, et al. (2002) Adiponectin reduces atherosclerosis in apolipoprotein E-deficient mice. Circulation 106(22):2767–70.

4. Lindberg S, et al. (2015) Adiponectin, type 2 diabetes and cardiovascular risk. Eur J Prev Cardiol 22(3):276–283.

5. Konter JM, et al. (2012) Adiponectin attenuates lipopolysaccharide-induced acute lung injury through suppression of endothelial cell activation. J Immunol 188(2):854–63.

6. Tao L, et al. (2007) Adiponectin Cardioprotection After Myocardial Ischemia/Reperfusion Involves the Reduction of Oxidative/Nitrative Stress. Circulation 115(11):1408–1416.

7. Vachharajani V, et al. (2012) Adiponectin-deficiency exaggerates sepsis-induced microvascular dysfunction in the mouse brain. Obesity (Silver Spring) 20(3):498–504.

8. Wang X, Buechler N, Yoza B, McCall C, Vachharajani V (2016) Adiponectin treatment attenuates inflammatory response during early sepsis in obese mice. J Inflamm Res Volume 9:167–174.

9. Yamauchi T, et al. (2007) Targeted disruption of AdipoR1 and AdipoR2 causes abrogation of adiponectin binding and metabolic actions. Nat Med 13(3):332–339.

10. Vasiliauskaité-Brooks I, et al. (2017) Structural insights into adiponectin receptors suggest ceramidase activity. Nature 544(7648):120–123.

11. Holland WLL, et al. (2013) An FGF21-adiponectin-ceramide axis controls energy expenditure and insulin action in mice. Cell Metab 17(5):790–7.

12. Xia JY, et al. (2015) Targeted Induction of Ceramide Degradation Leads to Improved Systemic Metabolism and Reduced Hepatic Steatosis. Cell Metab 22(2):266–278.

13. Wang Y, et al. (2014) Adiponectin inhibits tumor necrosis factor-α-induced vascular inflammatory response via caveolin-mediated ceramidase recruitment and activation. Circ Res 114(5):792–805.

14. Hug C, et al. (2004) T-cadherin is a receptor for hexameric and high-molecular-weight forms of Acrp30/adiponectin. Proc Natl Acad Sci U S A 101(28):10308–13.

15. Obata Y, et al. (2018) Adiponectin/T-cadherin system enhances exosome biogenesis and decreases cellular ceramides by exosomal release. JCI insight 3(8). doi:10.1172/jci.insight.99680.

16. Halberg N, et al. (2009) Systemic fate of the adipocyte-derived factor adiponectin. Diabetes 58(9):1961–70.

17. Yamauchi T, et al. (2003) Cloning of adiponectin receptors that mediate antidiabetic metabolic effects. Nature 423(6941):762–769.

18. Pajvani UB, et al. (2003) Structure-Function Studies of the Adipocyte-secreted Hormone Acrp30/Adiponectin. J Biol Chem 278(11):9073–9085.

19. Waki H, et al. (2003) Impaired Multimerization of Human Adiponectin Mutants Associated with Diabetes. J Biol Chem 278(41):40352–40363.

20. Liu Y, et al. (2007) Total and High Molecular Weight But Not Trimeric or Hexameric Forms of Adiponectin Correlate with Markers of the Metabolic Syndrome and Liver Injury in Thai Subjects. J Clin Endocrinol Metab 92(11):4313–4318.

21. Takemura Y, et al. (2007) Adiponectin modulates inflammatory reactions via calreticulin receptor-dependent clearance of early apoptotic bodies. J Clin Invest 117(2):375–86.

22. Galvan MD, Hulsebus H, Heitker T, Zeng E, Bohlson SS (2014) Complement protein C1q and adiponectin stimulate Mer tyrosine kinase-dependent engulfment of apoptotic cells through a shared pathway. J Innate Immun 6(6):780–92.

23. Kishore U, et al. (2004) C1q and tumor necrosis factor superfamily: modularity and versatility. Trends Immunol 25(10):551–61.

24. Ressl S, et al. (2015) Structures of C1q-like Proteins Reveal Unique Features among the C1q/TNF Superfamily. Structure 23(4):688–699.

25. Wesener DA, Dugan A, Kiessling LL (2017) Recognition of microbial glycans by soluble human lectins. Curr Opin Struct Biol 44:168–178.

26. Endo Y, Matsushita M, Fujita T (2011) The role of ficolins in the lectin pathway of innate immunity. Int J Biochem Cell Biol 43(5):705–712.

27. Seaton BA, et al. (2010) Review: Structural determinants of pattern recognition by lung collectins. Innate Immun 16(3):143–150.

28. Grundy SM (2016) Overnutrition, ectopic lipid and the metabolic syndrome. J Investig Med 64(6):1082–1086.

29. Chaurasia B, Summers SA (2015) Ceramides - Lipotoxic Inducers of Metabolic Disorders. Trends Endocrinol Metab 26(10):538–550.

30. Païdassi H, et al. (2008) C1q binds phosphatidylserine and likely acts as a multiligand-bridging molecule in apoptotic cell recognition. J Immunol 180(4):2329–38.

31. Min X, et al. (2012) Crystal structure of a single-chain trimer of human adiponectin globular domain. FEBS Lett 586(6):912–917.

32. Grosdidier A, Zoete V, Michielin O (2011) SwissDock, a protein-small molecule docking web service based on EADock DSS. Nucleic Acids Res 39(suppl):W270–W277.

33. Shapiro L, Scherer PE (1998) The crystal structure of a complement-1q family protein suggests an evolutionary link to tumor necrosis factor. Curr Biol 8(6):335–8.

34. Lu J, Kishore U (2017) C1 Complex: An Adaptable Proteolytic Module for Complement and Non-Complement Functions. Front Immunol 8:592.

35. Okamoto Y, et al. (2000) An Adipocyte-Derived Plasma Protein, Adiponectin, Adheres to Injured Vascular Walls. Horm Metab Res 32(02):47–50.

36. Almer G, et al. (2011) Globular domain of adiponectin: promising target molecule for detection of atherosclerotic lesions. Biologics 5:95–105.

37. Mori T, et al. (2015) Ultrastructural Localization of Adiponectin protein in Vasculature of Normal and Atherosclerotic mice. Sci Rep 4(1):4895.

38. Domingo-Almenara X, et al. (2018) XCMS-MRM and METLIN-MRM: a cloud library and public resource for targeted analysis of small molecules. Nat Methods 15(9):681–684.

39. Guijas C, et al. (2018) METLIN: A Technology Platform for Identifying Knowns and Unknowns. Anal Chem 90:3156–3164.

40. Sud M, et al. (2007) LMSD: LIPID MAPS structure database. Nucleic Acids Res 35(Database):D527–D532.

41. Tom Tang Y, et al. (2005) The complete complement of C1q-domain-containing proteins in Homo sapiens. Genomics 86(1):100–111.

42. Matsuda K (2017) Synapse organization and modulation via C1q family proteins and their receptors in the central nervous system. Neurosci Res 116:46–53.

43. Chavali VRM, Sommer JR, Petters RM, Ayyagari R (2010) Identification of a promoter for the human C1Q-tumor necrosis factor-related protein-5 gene associated with late-onset retinal degeneration. Invest Ophthalmol Vis Sci 51(11):5499–507.

44. Hirai H, et al. (2005) Cbln1 is essential for synaptic integrity and plasticity in the cerebellum. Nat Neurosci 8(11):1534–1541.

45. Elegheert J, et al. (2016) Structural basis for integration of GluD receptors within synaptic organizer complexes. Science 353(6296):295–9.

46. Rodriguez S, et al. (2016) Loss of CTRP1 disrupts glucose and lipid homeostasis. Am J Physiol Metab 311(4):E678–E697.

47. Martinelli DC, et al. (2016) Expression of C1ql3 in Discrete Neuronal Populations Controls Efferent Synapse Numbers and Diverse Behaviors. Neuron 91(5):1034–1051.

48. Zhang Y (2008) I-TASSER server for protein 3D structure prediction. BMC Bioinformatics 9(1):40.

49. Tu X, Palczewski K (2012) Crystal structure of the globular domain of C1QTNF5: Implications for late-onset retinal macular degeneration. J Struct Biol 180(3):439–46.

50. Wallis R, Mitchell DA, Schmid R, Schwaeble WJ, Keeble AH (2010) Paths reunited: Initiation of the classical and lectin pathways of complement activation. Immunobiology 215(1):1–11.

51. Peake PW, Shen Y, Walther A, Charlesworth JA (2008) Adiponectin binds C1q and activates the classical pathway of complement. Biochem Biophys Res Commun 367(3):560–565.

52. Peake P, Shen Y (2010) Factor H binds to the N-terminus of adiponectin and modulates complement activation. Biochem Biophys Res Commun 397(2):361–366.

53. Giles BM, Boackle SA (2013) Linking complement and anti-dsDNA antibodies in the pathogenesis of systemic lupus erythematosus. Immunol Res 55(1–3):10–21.

54. Boath A, et al. (2008) Regulation and traffic of ceramide 1-phosphate produced by ceramide kinase: comparative analysis to glucosylceramide and sphingomyelin. J Biol Chem 283(13):8517–26.

55. Buratta S, et al. (2017) Extracellular vesicles released by fibroblasts undergoing H-Ras induced senescence show changes in lipid profile. PLoS One 12(11):e0188840.

56. Jeon S-B, Yoon HJ, Park S-H, Kim I-H, Park EJ (2008) Sulfatide, a major lipid component of myelin sheath, activates inflammatory responses as an endogenous stimulator in brain-resident immune cells. J Immunol 181(11):8077–87.

57. van Zyl R, Gieselmann V, Eckhardt M (2010) Elevated sulfatide levels in neurons cause lethal audiogenic seizures in mice. J Neurochem 112(1):282–295.

58. Norton WT, Poduslo SE (1973) Myelination in rat brain: changes in myelin composition during brain maturation. J Neurochem 21(4):759–73.

59. Shaner RL, et al. (2009) Quantitative analysis of sphingolipids for lipidomics using triple quadrupole and quadrupole linear ion trap mass spectrometers. J Lipid Res 50(8):1692–707.

60. Folco EJ, Rocha VZ, López-Ilasaca M, Libby P (2009) Adiponectin inhibits pro-inflammatory signaling in human macrophages independent of interleukin-10. J Biol Chem 284(38):25569–75.

61. Phoonsawat W, Aoki-Yoshida A, Tsuruta T, Sonoyama K (2014) Adiponectin is partially associated with exosomes in mouse serum. Biochem Biophys Res Commun 448(3):261–266.

62. Crewe C, et al. (2018) An Endothelial-to-Adipocyte Extracellular Vesicle Axis Governed by Metabolic State. Cell 175(3):695–708.e13.

