## Supplemental Methods and Data for "Adiponectin and related C1q/TNF-related proteins bind selectively to anionic phospholipids and sphingolipids"

**SUPPLEMENTARY INFORMATION:**

EXTENDED MATERIALS AND METHODS

*Lipids*

1,2-dioleoyl-sn-glycero-3-phosphocholine (DOPC) (cat no. 850375), brain phosphatidylserine (PS) (cat no. 840032), C16 ceramide (Cer) (cat no. 860516), C16 ceramide-1-phosphate (Cer1P) (cat no. 860533P), Topfluor C11 ceramide (cat no. 810262), Topfluor C11 ceramide-1-phosphate (cat no 810270), brain sulfatide (ST) (cat no. 131305P), C16 cardiolipin (CL) (cat no. 710333P), C16 β-glucosylceramide (GlcCer) (cat no. 860539), NBD-PS (cat no. 810198), and rhodamine-phosphatidylethanolamine (PE) (cat no. 810150) were obtained from Avanti Polar Lipids. NBD-sulfatide (cat no. 1632-001) was obtained from Matreya LLC. Lipids were received in powdered form and resuspended in chloroform or 4:1 chloroform:methanol. Resuspended lipids were then stored in amber glass vials with Teflon coated caps at -20°C. Pipetting of stock solutions was performed with glass pipettes or Hamilton syringes (Sigma-Aldrich 26200-U).

*Western blot antibodies*

| Target | Type | Cat. No. | Vendor | Dilution |
|---|---|---|---|---|
| V5 | Mouse mAb | 46-0705 | Invitrogen | 1:5000 |
| 6x His tag | Rabbit | ab9108 | Abcam | 1:1000 |

*Plasmids and cloning*

pcDNA3.1 V5 His A was used as the vector backbone for all constructs in this study. Constructs were generated using Gibson assembly (NEB #E2611), restriction enzyme digest and ligation, or Q5 site-directed mutagenesis (NEB #E0554) following manufacture protocols. For Gibson assembly, full length Adipoq, CTRP1, and CTRP5 CDS were cloned from mouse visceral white adipose tissue cDNA, and Cbln1 CDS from mouse hindbrain cDNA, using Phusion HF DNA polymerase (NEB M0530) and Gibson assembly primers designed using the NEBuilder online tool. pcDNA3.1 V5 His A vector was either linearized with by PCR with Phusion and Gibson primers, or cut using Xho1 (NEB #R0146) and/or BstBI (NEB #R0519). CTRP13 was initially cloned in a pSECTAG vector behind an Igκ leader sequence, then transferred to a pcDNA3.1 V5 His backbone with the Igκ leader sequence intact. Site directed mutagenesis was performed using the NEB Q5 site directed mutagenesis kit (NEB E0554); primers for specific locations and mutations were designed using the NEBaseChanger online tool. All final plasmids were sequence-verified through the entire coding region.

*Cloning Primers*

| Target/Product | Forward (5' to 3') | Reverse (5' to 3') |
|---|---|---|
| ***<u>Gibson assembly (and restriction sites)</u>*** | | |
| mAdipoq | agggcccttcgaaccaagctggctagttaATG CTACTGTTGCAAGCTCTC | tagggataggcttaccGTTGGTAT CATGGTAGAGAAGAAAG |
| pcDNA3.1 (w/ Adipoq) | GGTAAGCCTATCCCTAACCC | taactagccagcttggTTCGAAGG GCCCTCTAGAC |
| mRbp4 | agagggcccttcgaaATGGATCCTCTG GGCTGG | tagggataggcttaccCAAACTGT TTCTGGAGGGC |
| pcDNA3.1 (w/ Rbp4) | GGTAAGCCTATCCCTAACCC | TTCGAAGGGCCCTCTAGAC |
| mCTRP5 (BstBI) | ctcgagtctagagggcccttATGAGGCCA CTTCTTGCC | gttagggataggcttacctttAGCGA AGACTGGGGAGCTG |
| mCTRP1(Xho1/BstBI) | cacagtggcggccgcATGGGCTCCTGT GCACAG | gggataggcttaccttgGGGCTCA GAGGCTGGCTT |
| mCbln1 (BstBI) | gtctagagggcccttATGCTGGGCGTC GTGGAG | gggataggcttaccttgGAGGGGA AACACGAGGAATCC |

| *Site directed mutagenesis* | | |
|---|---|---|
| mAdipoq collΔ | GCTTATGTGTATCGCTCAGC | TGCCATCCAACCTGCACA |
| mAdipoq C1qΔ | GGTAAGCCTATCCCTAACC | GGCTTCTCCAGGCTCTCC |
| mAdipoq H224A | GGATGGGGACgcaAATGGACTCT ATGCAGATAAC | CCATACACCTGGAGCCAG |
| mAdipoq R134A | TGTACCCATTgccTTTACTAAGATC TTCTACAACC | TTGGGAACAGTGACGCGG |
| mAdipoq C39S | CAAGGGAACTagtGCAGGTTGGAT | GGTGGAGGGACCAAAGCA |
| mAdipoq G87R | AGGAGATGTTcgtATGACAGGAGC TG | GTCTCACCCTTAGGACCA |

*Protein expression*

The Expi293 mammalian cell expression system (ThermoFisher) was used for expression of recombinant adiponectin and CTRP family members. Cells were cultured in Expi293 expression medium on a shaking platform in a tissue culture incubator at standard tissue culture conditions (37°C, 5% $CO_2$, 95% humidity). Before transfection, cells were resuspended to a density of 1 x $10^6$ cells/mL. Plasmid DNA (10 µg/mL) and linear PEI (30 µg/mL) (MW 25000, PolySciences) were then mixed at a 1:3 ratio (w/w) in PBS and incubated at room temperature for 20 min to allow encapsulation. The mixture was then added to cells (1 µg DNA per $10^6$ cells), and incubated for 5 days in standard cell culture conditions.

*Protein isolation by Ni-NTA affinity chromatography*

Expressed protein was first harvested from Expi293 cell supernatant and filtered through a 0.22 µm-pore filter (Corning 431118) to remove residual cellular debris. Ni-NTA agarose beads (Qiagen 30210) were then added to the supernatant, plus 10 mM imidazole to reduce non-specific bead binding, and incubated overnight in a 4°C rotator. Roughly 1 mL of packed beads was used per 100 mL supernatant. Beads were then washed in batch three times in 20 mM imidazole and transferred to a 5 mL disposable column (Qiagen 1000333). Protein was then eluted from beads using 250 mM imidazole in HNC isolation buffer (25 mM HEPES (AmericanBio AB06021-00100), 150 mM NaCl, 1 mM $CaCl_2$, pH 7.4). All steps performed on ice or at 4°C. Elution fractions were assessed by Western blot and silver stain, along with samples of input, flow through, and wash steps. After analysis, fractions with protein were collected and concentrated using Amicon-Ultra 3- or 10-kDa molecular weight cut-off centrifugal spin filters (Millipore). Protein samples were then washed three times with HNC buffer to remove imidazole, and the A280 of the final isolate was measured by nanodrop. Protein was then kept at 4°C until use or snap frozen in liquid nitrogen and kept at -80°C for longer storage.

*Gel filtration*

In some cases, isolated protein was analyzed by size exclusion chromatography (Superdex 200 Increase 10/300) using HNC buffer. Eluted protein was concentrated and either used directly or aliquoted and flash-frozen for later use in biochemical assays.

*SDS-PAGE / Western blot*

Samples for Western blot were mixed with 5x SDS sample buffer and boiled for 5 min before loading and running on a 4-15% Mini-Protean TGX precast protein gel (Bio-Rad 456-1086). Protein bands were then transferred to a 0.45 μm pore PVDF membrane (Bio-Rad IMMUN-BLOT, Millipore Immobilon-P IPVH00010) using a semi-dry Trans-Blot Turbo Transfer system (Bio-Rad). Membranes were blocked with 5% milk in Tris-buffered saline with 0.1% tween (TBST), or 5% BSA in TBST for phospho-protein blots, for at least 1 hr at room temperature. Blots were then probed with primary antibody overnight at 4°C, washed 3 times with TBST, then incubated with goat anti-mouse or goat anti-rabbit peroxidase-conjugated secondary antibody (Jackson ImmunoResearch 111-035-045, 115-035-062) for 40 min before adding enhanced chemiluminescence (ECL) reagent (ThermoFisher 32106). Bands were then visualized by exposing blots to film (Denville E3018) under darkroom illumination and developing on a Kodak X-OMAT 200A Processor. For some blots with weak signal, femto ECL reagent (ThermoFisher 34095) was used and chemiluminescent signal was visualized on a GelDoc imager (Bio-Rad). Films were then scanned and quantified using FIJI software.

*SDS-PAGE / Silver stain*

Samples for silver stain were mixed with 5x SDS sample buffer and boiled for 5 min before loading and running on a NuPage 4-12% Bis-Tris polyacrylamide gel (Invitrogen NP0323). Gels were then excised and incubated overnight in fixer (40% methanol, 13.5% formalin in water). Following fixation, the gels were pretreated with 0.02% (w/v) sodium thiosulfate, stained with 1 mg/mL $AgNO_3$ for 15 minutes, and treated with developer (3% (w/v) sodium carbonate, 0.004% (w/v) sodium thiosulfate, and 0.05% (v/v) formalin in water) for 5-10 minutes or until clear bands were seen. Reaction was stopped with 20% (w/v) acetic acid and the resulting gel was placed on a light box and photographed.

*Dot blot*

Dot blots were either obtained commercially from Echelon Biosciences and Avanti Polar Lipids, or made from candidate lipids spotted on nitrocellulose strips. In brief, candidate lipids were dissolved in 4:1 chloroform:methanol solvent at a concentration of 1 mg/mL. A Hamiltonian syringe was then used to spot 2 uL of the lipid solution on a dry, standard 0.45 μm pore nitrocellulose membrane (Millipore). The membranes were allowed to dry overnight at 4°C before use. Dot blots were then blocked using 3% fatty acid-free BSA in TBST, and incubated with tagged, expressed protein in unpurified cell culture supernatant overnight. In some cases, a positive anti-V5 antibody was spotted onto the membrane to use a positive control and to approximate abundance of tagged protein. Blots were subsequently washed and probed with rabbit anti-6x His primary antibody (1:1000), followed by goat anti-rabbit peroxidase-conjugated secondary antibody (Jackson ImmunoResearch). Blots were developed using ECL reagent (ThermoFisher 32016), imaged using a GelDoc imager (Bio-rad) using the chemiluminescence protocol, and analyzed using FIJI software.

*Liposome synthesis*

Lipids in solvent were mixed at defined molar ratios in an ultra-clear microcentrifuge tube (Axygen MCT-175-C), and evaporated under nitrogen stream. After evaporation, the lipid film was dried under vacuum for 1 hr at room temperature in a VWR 1410 vacuum oven, then rehydrated using HNC buffer, or 0.75M sucrose in HNC for sucrose-loaded liposomes. For rehydration, buffer was added to the lipid film to form a 5 mM solution and subjected to 10 freeze-thaw cycles with occasional vortexing between cycles. For creation of liposomes with defined size, the mixture was then passed 11 times through a manual Avestin lipid extruder equipped with either a 100 nm or 1000 nm filter. Sucrose loaded liposomes were then diluted with HNC, pelleted by centrifugation (15000 rpm x 15 min at 4°C), and resuspended in the original volume of HNC. Synthesized liposomes were then stored at 4°C until use, or kept at -80°C for longer storage.

*Liposome pull-down assay*

1000 nm sucrose-containing liposomes of varying lipid composition were generated using freeze-thaw cycling and extrusion. In addition to the specified amount of target lipid, 1% rhodamine-PE was added to help visualize the liposomes. After liposome synthesis, 5 µL liposomes (5 mM stock) were combined with 10 µL of 50 mg/mL fatty acid-free BSA and 25 µL HNC in Eppendorf tubes. The mixture was incubated for 5 minutes at room temperature to “block” the liposomes, then 10 µL of purified recombinant protein at 50 µg/mL was added to the mixture to make a final concentration of 10 µg/mL in a total assay volume of 50 µL. Volumes of liposome or protein were increased as needed if concentrations of either stock solutions were too low, and buffer amounts were decreased commensurately to maintain a total volume of 50 µL. Reduced protein, when used, was prepared by adding β-mercaptoethanol to a final concentration of 2% (0.28 mM) to the 10 µg/mL recombinant protein stock before addition to lipids. Final mixtures of protein and liposome were then incubated at 37°C for 1 hr to allow binding to occur. After binding, liposomes were diluted with 500µl of HNC and spun down at 15000 rpm x 15 min. Supernatants were aspirated (and saved), then the pellets were washed an additional time with 500 µL HNC. Finally, pellets were resuspended in 25 µL HNC and added to a black opaque low-adsorption 96-well assay plate (Corning 3991) containing 3 µL 10% Triton X-100 to measure rhodamine fluorescence. After measurement, samples were denatured with 5x SDS sample buffer, boiled, and analyzed by Western blot. For this assay, samples were denatured in non-reducing conditions unless specified otherwise.

*Lipid transfer assay*

Labeled (donor) and unlabeled (acceptor) 100 nm or 1000 nm liposomes were generated as described using freeze-thaw cycling and extrusion. Donor liposomes contained equimolar amounts of NBD- or Topfluor labeled ligand and rhodamine-PE quencher. Acceptor liposomes were 100% phosphatidylcholine. After liposome synthesis, candidate protein isolates were plated at 10 µg/mL in a black opaque low-adsorption 96-well assay plate (Corning 3991) in HNC buffer, and 1.5 µl donor and acceptor liposomes (5 mM stock) were added to a final concentration of 150 µM each in a total assay volume of 50 µl. A SpectraMax M5 plate reader (Molecular Devices) was then used to monitor fluorescence at excitation/emission 480/530 every 10 seconds for 1.5 hrs to assess loss of FRET quenching of NBD or Topfluor by rhodamine. For some experiments, at the completion of the timecourse, the liposomes were lysed with 10 µl *of 10% Triton X-100* to measure the maximum achievable fluorescence, to which the timecourse was normalized.

*Protein-bound lipid analysis by liquid chromatography-mass spectrometry (LC-MS) and tandem mass spectrometry (MS/MS)*

Expressed proteins were isolated as described from transfected cell supernatant and, after elution and buffer exchange, concentrated to 80-100 ul using an Amicon-Ultra 3 kDa MWCO spin filter. Samples were then collected into a polypropylene microcentrifuge tube, with atmosphere switched to nitrogen to prevent lipid oxidation. Individual tubes were snap frozen in liquid nitrogen and stored at -80°C until shipping. Subsequent LC-MS and MS/MS analysis was performed by the Scripps Center for Metabolomics and Mass Spectrometry. Upon receipt, samples on dry ice were thawed and extracted with 400ul of ice-cold 50:50 methanol:acetonitrile (ACN). Lipid extracts were then analyzed by LC-MS using a Bruker Impact II Q-TOF coupled to an Agilent 1200 LC stack. Mobile phases consisted of A = 95:5 $H_2O$:ACN/0.1% formic acid, and B = 9:1 IPA:ACN/0.1% formic acid. Column used was an Agilent 300A SB-C18 1.0mm x 150mm running at 50ul/min. Positive ions were analyzed using XCMS software. MS/MS analysis was performed on m/z = 870.74. The resulting MS/MS spectral peaks were then searched against various databases in METLIN and LipidMaps. The prediction for sulfatide was found using the glycerophospholipid search tool in LipidMaps (http://www.lipidmaps.org/tools/ms/glycosylcer_gen.html).

*Low density lipoprotein flotation assay*

7.5 μl of human LDL solution (Sigma-Aldrich L7914) was mixed with 10 μl fatty-acid free BSA at 50 μg/mL, 17.5 μl HNC buffer, 5 μl Sudan black (10 mg/ml in ethanol, Sigma-Aldrich 199664), and 10 μl recombinant protein at 50 μg/mL as in liposome pulldown assays. Reduced protein, when used, was prepared by adding β-mercaptoethanol to a final concentration of 2% (28 mM) to the 10 μg/mL recombinant protein stock before addition to lipids. Lipid-protein mixtures were incubated for 30 minutes at 37°C to allow lipid binding. After the incubation period, the mixture was mixed with 280 µl of OptiPrep (a 60% iodixanol solution with density 1.32 g/mL, Stem Cell 07820) to form a final 80% OptiPrep mixture (density 1.25 g/mL), which was added to the bottom of a 2.2 mL thin-walled UltraClear ultracentrifuge tube (Beckman Coulter 347356). This was then overlaid with 330 µl 70% OptiPrep in HNC (density 1.22 g/mL), 670 µl 36% OptiPrep in HNC (density 1.1 g/mL), and 670 ul HNC buffer (density 1.0 g/mL). Gradients, prepared in pairs, were then loaded into a TLS-55 ultracentrifuge rotor and spun at 50000 rpm x 20 hrs at 4°C with no brake on an Optima TL benchtop ultracentrifuge (Beckman Coulter). Upon completion of ultracentrifugation, tubes were retrieved and fractions were collected by puncture from the bottom of the tube. Briefly tubes were punctured with a 20G needle and roughly 100 µl fractions were withdrawn into a syringe. Sequential fractions were plated into a transparent 96-well plate, and absorbance of Sudan black was read at 600 nm using a SpectraMax M5 plate reader (Molecular Devices). Selected fractions were then collected into 5x SDS sample buffer for detection of recombinant protein by Western blot. In general, non-reducing conditions were used for this analysis.

*Plasma collection and lipidomics*

8-10-week old male WT C57B6/J (The Jackson Laboratory Stock No. 000644) and Adipoq -/- mice (B6;129-Adipoqtm1Chan/J, The Jackson Laboratory Stock No. 008195) were housed in an SPF-facility with regulated 12 hr light-dark cycles with *ad libitum* access to standard chow and water. Whole blood was harvested by retro-orbital bleeding, and plasma was isolated using lithium-heparin coated plasma separator tubes (BD 365985) and stored at -20°C. Lipidomics was performed by the West Coast Metabolomics Center at UC Davis. In brief, extraction of samples was performed in MTBE with addition of internal standards, followed by ultra-high-pressure liquid chromatography on a Waters CSH column, interfaced to a QTOF mass spectrometer (high resolution, accurate mass), with a total run time of 15 minutes. Data were collected in both positive and negative ion mode, and analyzed using MassHunter (Agilent).

Approximately 400 lipids can be identified from plasma, with additional unknowns. The method is highly stable and has been validated on large datasets (>8,000 samples) collected over long time periods (> 1 year). Counts of each lipid species for each sample was normalized to the counts of the closest corresponding internal standard to obtain normalized peak height. To depict differences over various lipid classes, normalized peak heights for each lipid species within a class were summed, and each sum was then normalized to the average of the sums over the WT samples to assess relative fold change.

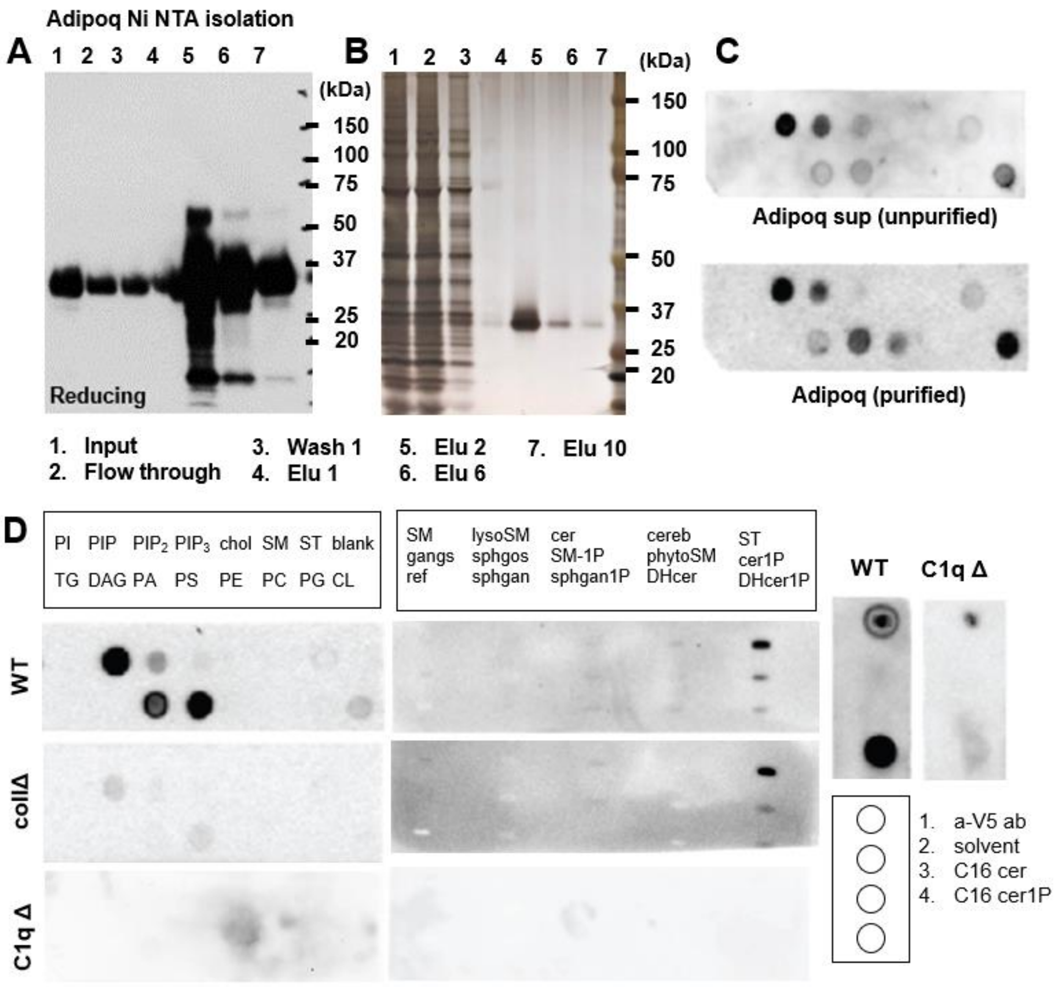


**Supplementary figure S1. Purified adiponectin binds specifically to lipids in lipid strips.**

(A-B) Ni NTA isolation of recombinant full-length murine adiponectin cloned into pcDNA3.1 v5 His plasmid and expressed in Expi293 cells.

(A) Western blot of isolation fractions probed using anti-V5 antibody showing expression and elution of protein. Though samples run in reducing buffer, residual incompletely reduced dimer (~60 kDa) can still be seen.

(B) Silver stain showing purity of eluted protein. Samples run in reducing conditions.

(C) Echelon lipid strips probed with unpurified Adipoq-V5 His transfected supernatant (left) and Adipoq purified protein isolate (right) showing comparable binding.

(D) Representative lipid strips (Echelon left, Avanti middle, in-house right) showing binding of WT adiponectin and mutants with deletion of collagenous domain (collΔ) or of C1q head domain (C1qΔ).

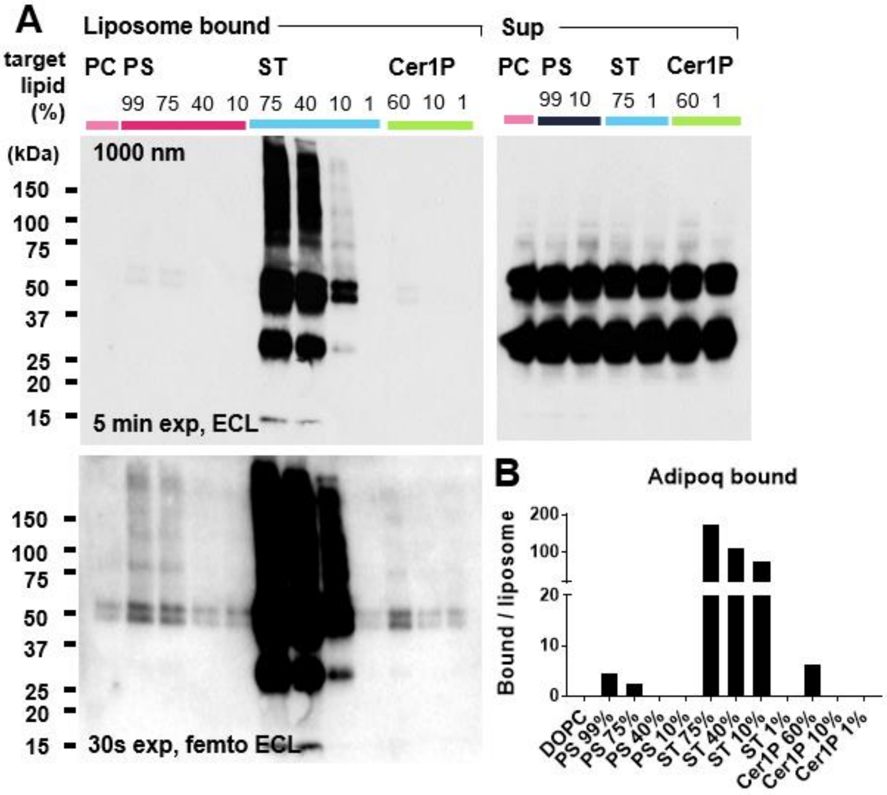


**Supplementary Figure S2. Adiponectin lipid binding requires a relatively high density of target lipids.**

(A) Western blot of adiponectin bound to liposomes of varying target lipid composition in pulldown assay. Adiponectin in pellet fractions shown as liposome bound; excess adiponectin in the supernatant (diluted 1:20) for the various conditions shown at right for reference. Percentage of target lipids in liposomes indicated above each lane. Size of liposomes (1000 nm) shown in upper left corner. Second exposure with femto ECL (middle) shows better visualization of weaker bands.

(B) Densiometric quantification of Western blot above. Band densities normalized to total rhodamine fluorescence of liposome pellet. n = 1 experiment shown.

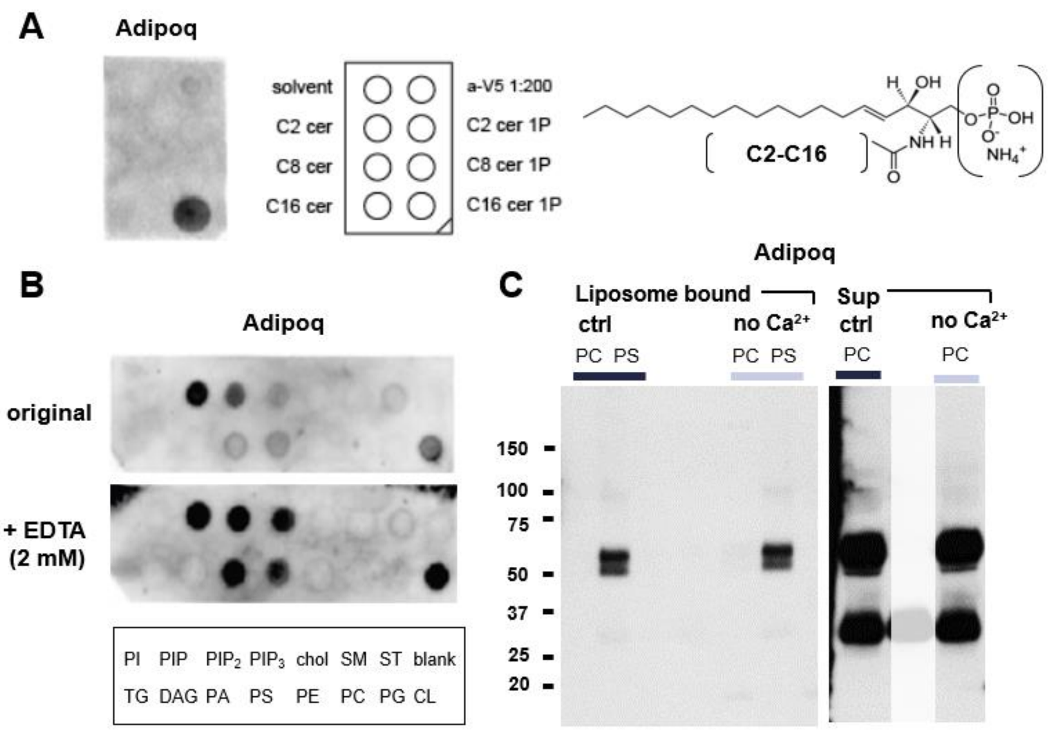


**Supplementary figure S3. Tail length of ceramide fatty acyl group, but not $Ca^{2+}$, affects adiponectin binding to lipids.**

(A) Binding of adiponectin to ceramides and ceramide-1-phosphates of various fatty acyl tail lengths spotted on lipid strips. Tail length of fatty acyl chain of ceramide indicated by C designation (C2 = 2 carbons, C8 = 8 carbons, C16 = 16 carbons). Anti-V5 antibody (a-V5 1:200 dilution) spotted as a positive control.

(B) Binding of adiponectin in to lipids in lipid strips in the presence or absence of $Ca^{2+}$ chelator EDTA.

(C) Liposome pull-down of adiponectin with or without removal of $Ca^{2+}$ from liposome and adiponectin buffers by buffer exchange with calcium-free HEPES-buffered saline. PC = 99% DOPC, PS = 99% phosphatidylserine.

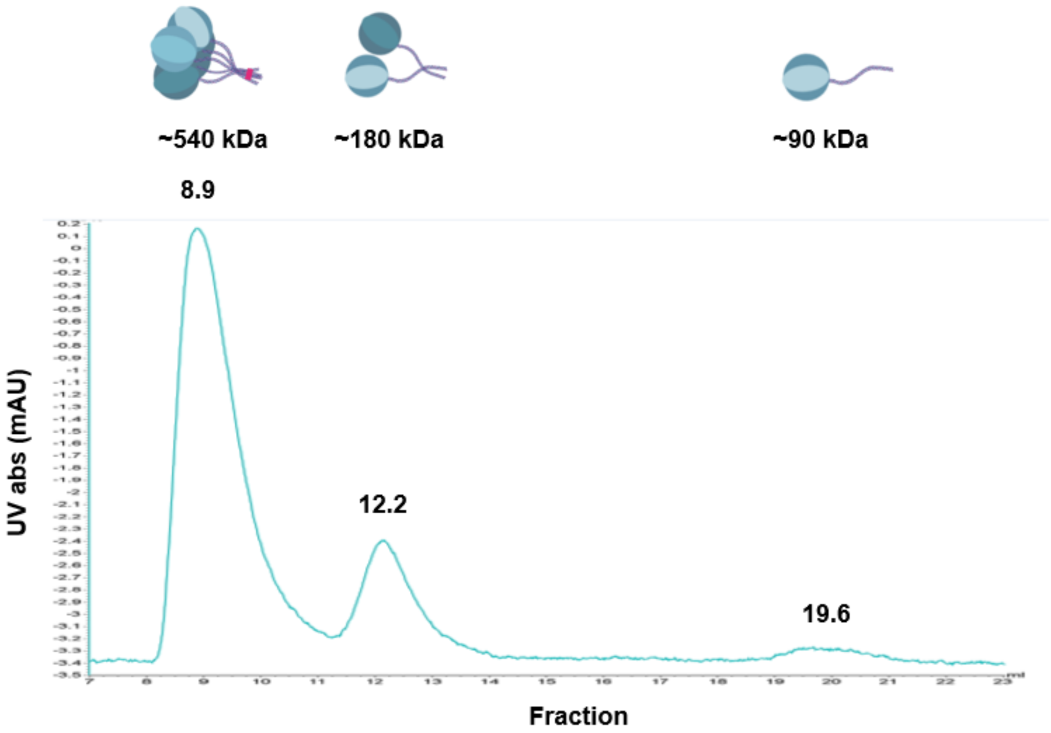


**Supplementary figure S4. Recombinant adiponectin expressed in Expi293 cells forms higher molecular weight oligomers.** UV absorbance profile of freshly isolated recombinant adiponectin expressed from Expi293 cells subjected to gel-filtration chromatography on a Superose 200 column (flow rate 0.5 mL/min). Fraction numbers of protein elution peaks annotated above each peak. Elution time of peaks consistent with the molecular weight of LMW trimers, MMW hexamers (dimers of trimers), and HMW oligomers (5-6-mers of trimers). Representative of n = 2 experiments.

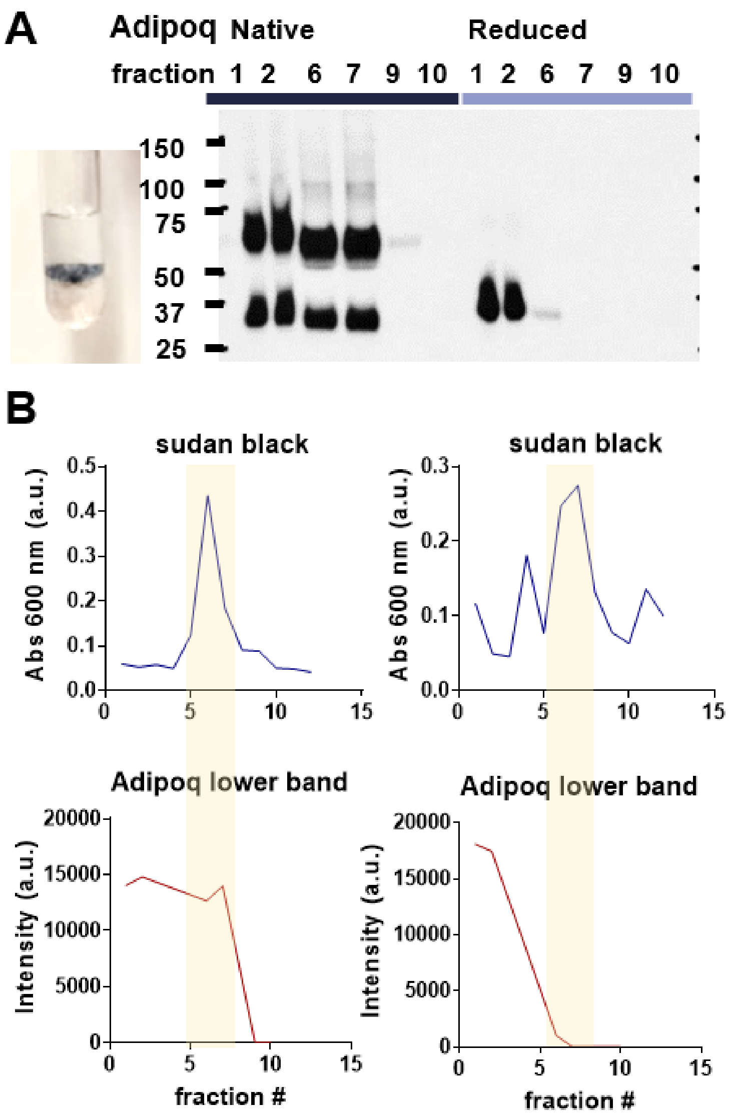


**Supplemental Figure S5.** Reduction of adiponectin oligomers abrogates binding to LDL. (A) Representative Western blots of adiponectin protein in sequentially eluted fractions of iodixanol gradients. Fraction number written above each lane. Inset photographs show banding pattern of LDL, stained with Sudan black, within the iodixanol gradient. (B) Quantification of Sudan black absorbance in sequential fractions (upper) and adiponectin protein in Western blots (lower). Yellow bars highlight Sudan black peaks corresponding to the location of LDL in the gradient.

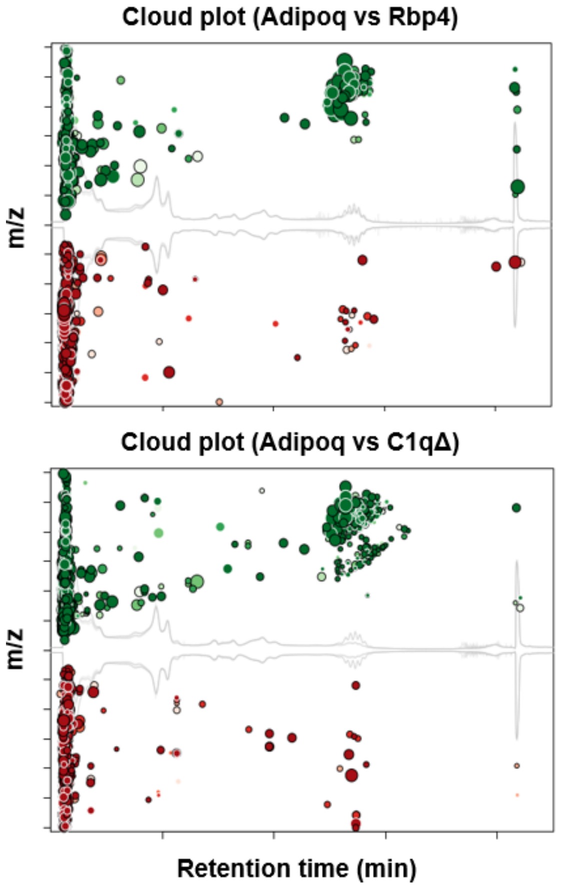


**Supplementary Figure S6. Untargeted LC-MS shows lipid peaks uniquely bound to adiponectin via the C1q head domain**.

Cloud plots showing global differences in lipid ligands found in solvent-extracted isolated protein by LC-MS. Column retention time plotted on the x-axis; m/z plotted on the y-axis. Dot size corresponds to peak height; darkness to p-value. Green dots are adiponectin-bound ligands while red dots are ligands bound to Rbp4 or C1qΔ controls.

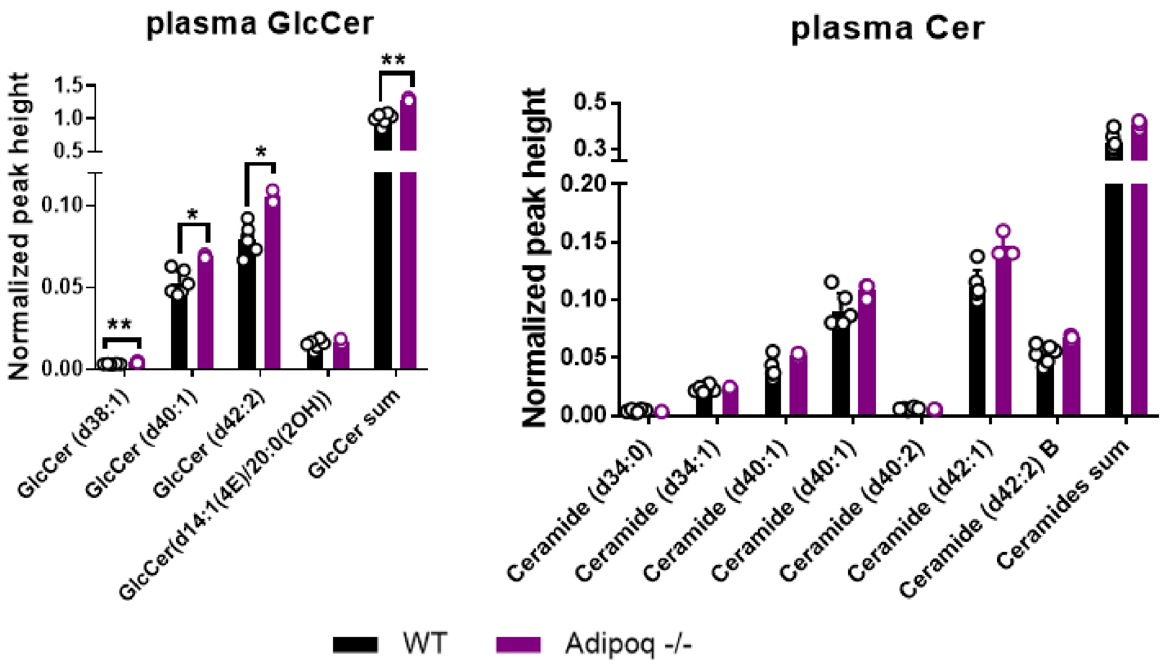


**Supplementary Figure S7. Shotgun lipidomics analysis of plasma of WT and Adipoq KO mice.** Peak intensity reads, normalized to that of the corresponding internal standard, for glucosylceramide and ceramide species, calculated and graphed individually as fold change relative to the WT average for that lipid (above). Error bars reflect S.D. from the mean. Open circles represent values from individual mice. ** $p < 0.01$, * $p < 0.05$. Unmarked bars were found to be not significantly different from WT.

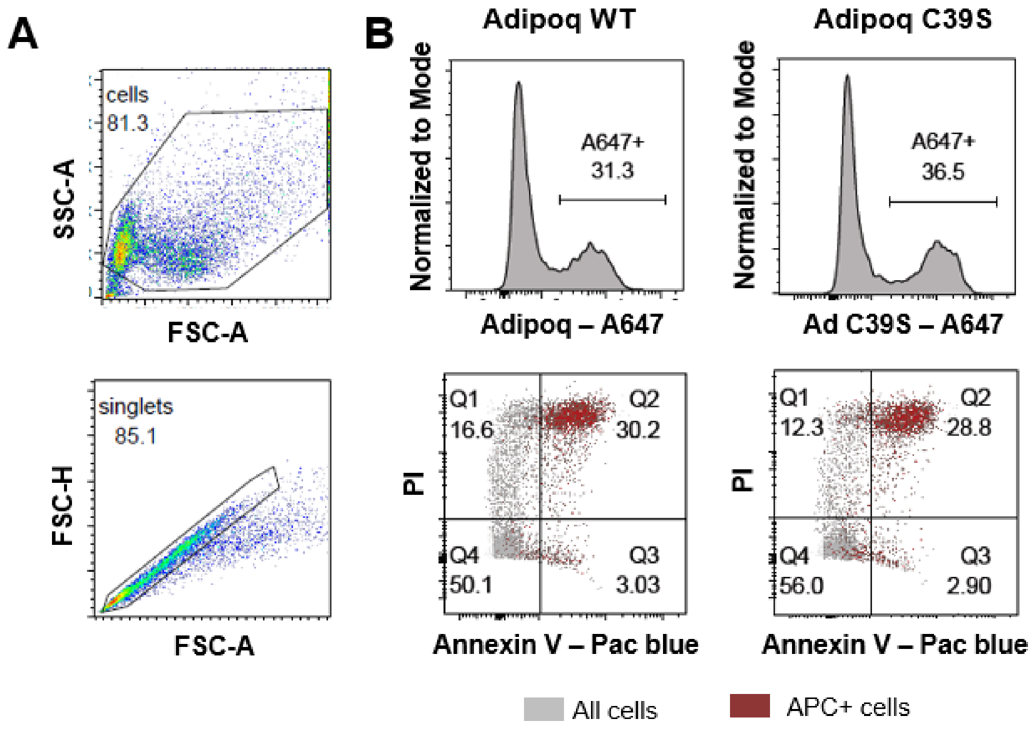


**Supplementary Figure S8. Adiponectin binds to dead and apoptotic cells, but not in an oligomerization-dependent manner.**

(A) Gating strategy for Expi293 cells used for staining with WT / C39S adiponectin, Annexin V, and PI. A wide gate was used to capture live and dead cells while excluding small debris (above) which was further refined using a singlet gate (below).

(B) (Top) Representative histograms of Expi293 singlets showing population level staining with WT / C39S adiponectin conjugated to Alexa 647. (Bottom) Annexin V vs PI staining of protein-bound A647+ cells (red) overlaid with that of total stained cells (grey) for both WT and C39S adiponectin constructs. Quadrant percentages displayed reflect that of total stained cells in grey.

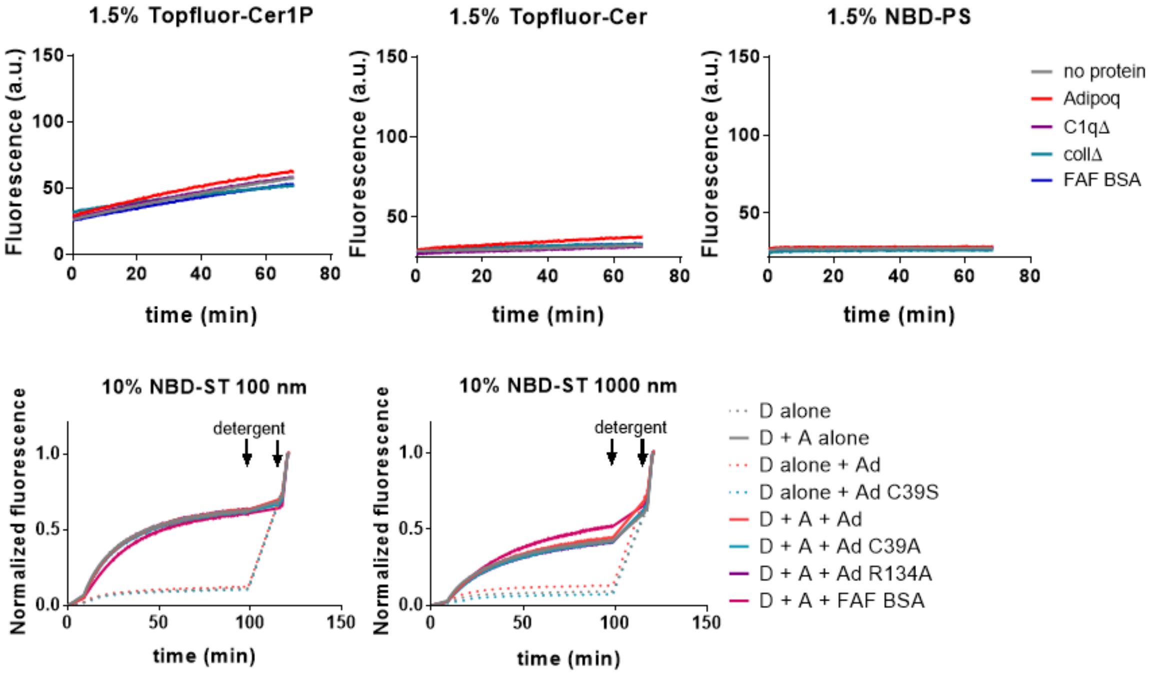


**Supplementary Figure S9. Adiponectin is not able to transfer individual target lipids between membranes.** Liposome transfer assay monitoring signal due to loss of FRET quenching upon transfer of various TopFluor or NBD tagged fluorescent target lipid from quenched donor liposomes to free acceptor liposomes. Ligand targets specified in the title of each graph. In upper panels, absolute TopFluor / NBD fluorescence is plotted on the y-axis. In lower panels, fluorescence signal was normalized to maximum fluorescence achieved after addition of detergents to de-quench all liposomes. D = donor liposome, A = acceptor liposome, Ad = adiponectin, FAF BSA = fatty acid free bovine serum albumin, Cer = ceramide, Cer1P = ceramide-1-phosphate, PS = phosphatidylserine, ST = sulfatide.

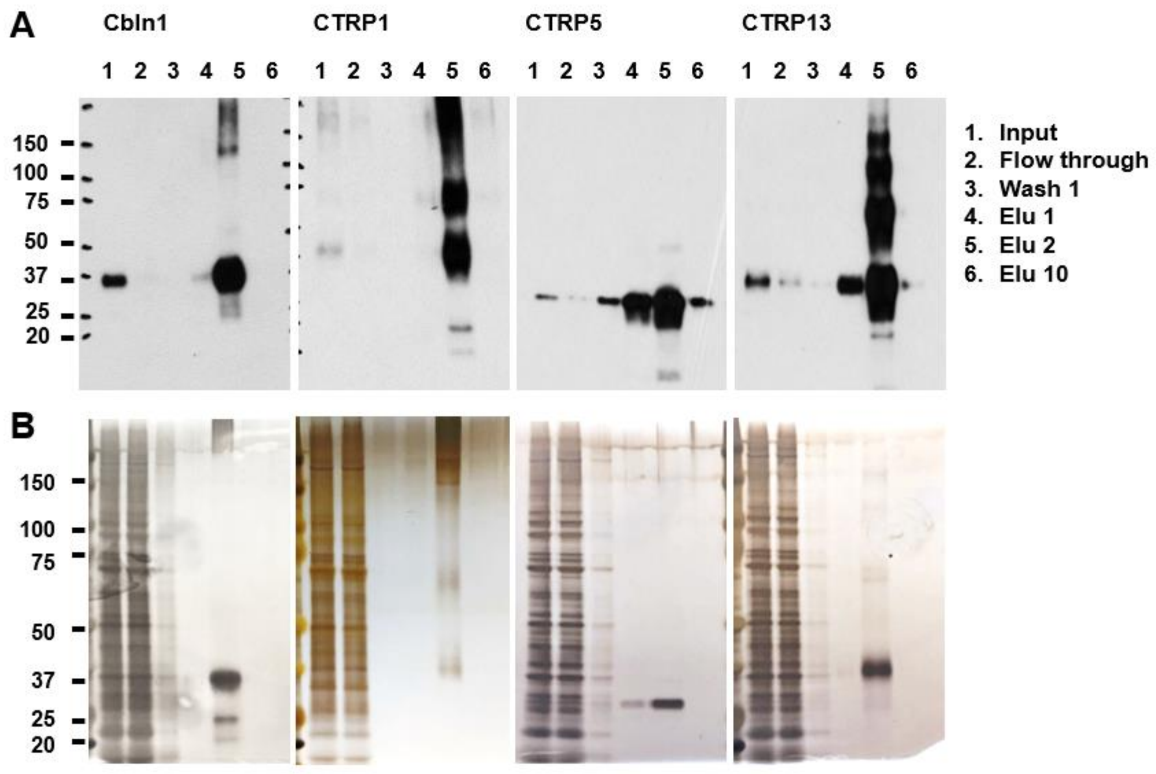


**Supplementary figure S10. Expression and isolation of recombinant murine CTRP family member proteins.**

(A) Western blot of isolation fractions probed using anti-V5 antibody showing expression and elution of various CTRP family proteins. Samples run in reducing conditions.

(B) Silver stain of input and eluted protein showing reasonable purity of protein of interest.

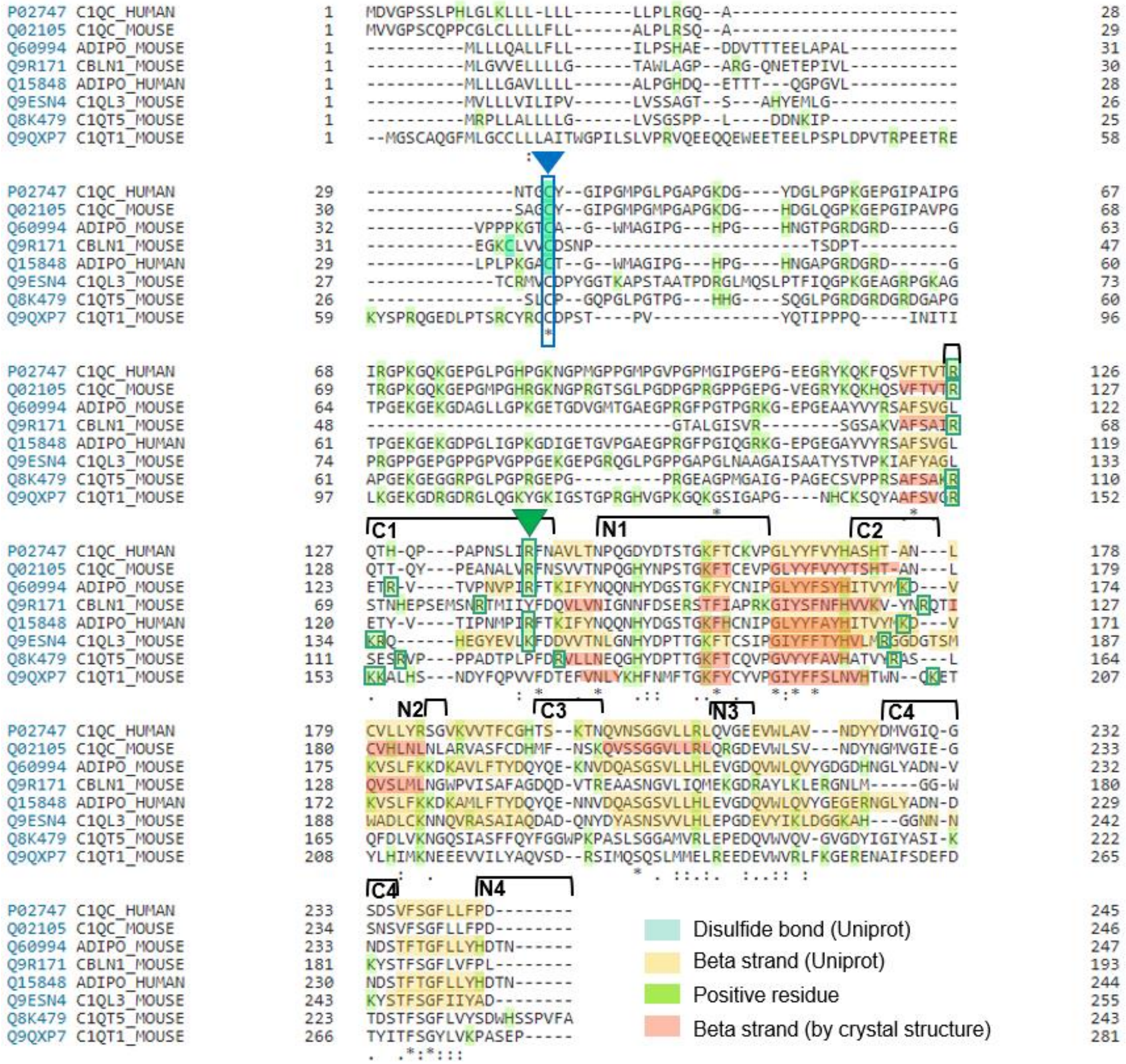


**Supplementary Figure S11. Sequence alignment of mouse and human adiponectin, mouse and human C1qC subunit, and mouse Cbln1, CTRP1, CTRP5, and CTRP13.** Amino acid sequences for each protein aligned using Uniprot Align tool. C39 residue of adiponectin relevant for oligomerization, and analogous conserved cysteines in other orthologues and paralogues, indicated in blue with blue arrow. Residues putatively involved in beta sheets, as annotated by Uniprot, shaded in yellow. Residues found involved in beta sheets by inspection of available crystal structures in PyMol or obtained by I-TASSER threading algorithms annotated in orange. Intervening solvent exposed loops marked by black brackets, as well as annotated by whether they appear on the N- or C-terminal end of the globular domain. Positive residues highlighted in green. Lysines and arginines in solvent exposed loops on C terminus of protein potentially involved in selectivity for negatively charged lipid ligands marked by green squares. R134 residue of adiponectin targeted in site directed mutagenesis marked by green arrow.
